# The non-coding facet of *Pink1* mRNA regulates mitochondria homeostasis

**DOI:** 10.64898/2026.01.05.697839

**Authors:** Xiaofen Wu, Yuying Liu, Ruimiao Ma, Hongye Wang, Ruqi Fan, Hui Wang, Weili Yang, Xiaojiang Li, Guoliang Xu, Hong Cheng, Ping Hu

## Abstract

Coding and non-coding RNAs have traditionally been viewed as functionally distinct entities. Here, we found that the down-regulation of *Pink1* due to *Tet2* knockout (KO) in muscle cells resulted in mitochondria dysfunction and muscle fatigue. Surprisingly, both *Pink1* mRNA alone and untranslatable ATG-deficient *Pink1* RNA were sufficient to rescue the mitochondria defects in *Tet2* KO cells. PINK1 protein was fast degraded shortly after translation and dispensable for functions of *Pink1* mRNA in muscle cells. Moreover, a portion of *Pink1* mRNA was located in mitochondria and interacted directly with mitochondria inner membrane located protein YME1L1, promoting its proteolytic cleavage of Opa1, a master regulator of mitochondria fusion, to generate the mature protein form. These findings uncovered a bona fide non-coding role for a canonical mRNA, suggesting that translation independent functions could be present in mRNAs with coding potentials, blurring the conventional dichotomy between coding and non-coding RNAs.

## INTRODUCTION

DNA methylation dynamics play critical roles in epigenetic regulations of many cellular processes. The Ten-eleven translocation (Tet) family DNA dioxygenases catalyze the stepwise oxidation of 5-methylcytosine (5mC) to 5-hydroxymethylcytosine (5hmC), and subsequently to 5-formylcytosine (5fC) and 5-carboxylcytosine (5caC), thereby facilitating active DNA demethylation (*1, 2*). These modifications play essential roles in the control of gene expression across diverse biological processes (*3–7*). Tet2 has been shown to regulate myogenesis by reducing DNA methylation near MyoD binding sites, enhancing MyoD binding affinity at these hypomethylated sites and activating myogenesis genes (*8*). However, how DNA methylation dynamics influences mitochondrial function in muscle cells remains to be fully elucidated.

Skeletal muscle is a highly energy-demanding tissue essential for force generation and locomotion. Muscle fatigue, characterized by a decline in contractile force and endurance, significantly impairs mobility and overall muscle function (*9, 10*). Muscle weakness and fatigue are common symptoms in aging and various diseases, including sarcopenia, chronic obstructive pulmonary disease (COPD), cancer, and chronic inflammation (*11*). A variety of factors, such as limitations in energy supply, impaired calcium signaling, the accumulation of lactate and other metabolic byproducts have been attributed to muscle fatigue (*10, 12, 13*). Mitochondria dysfunction is widely recognized as a central contributor to muscle fatigue and impaired muscle function (*14–18*). Given their pivotal role in cellular energy production and metabolism, maintaining mitochondrial integrity is crucial for optimal muscle performance. *Pink1* (*PTEN induced kinase 1*), a gene associated with early-onset Parkinson’s disease, has been shown to play important roles in mitochondrial quality control and cellular stress responses in brain and other tissues (*19–22*). However, the mechanisms regulating *Pink1* expression and its functions in skeletal muscle remain to be poorly understood. Long non-coding RNAs (lncRNAs) are conventionally defined as transcripts longer than 200 nucleotides lacking protein coding capability (*23*). They have been widely reported to play critical roles in myogenesis, muscle atrophy, and metabolism (*24–27*). These functions are achieved through diverse mechanisms, including direct interactions with key myogenic transcription factors such as MyoD, YY1, and Dnmt (*28–31*), modulation of chromatin accessibility and promoter activity (*32–35*), regulation of mRNA translation (*36*), and sequestration of microRNAs (miRNAs) as competing sponges (*37–42*). For example, *Linc-RAM* activates the transcription of *Myog* and *Myh3* transcription by interacting with MyoD (*28, 29*). Enhancer RNAs (eRNAs), such as *MUNC* and *^CE^RNA*, promote myogenic gene expression by facilitating chromatin remodeling and recruitment of key transcription factors (*43, 44*). By contrast, messenger RNAs (mRNAs) have traditionally been viewed as passive intermediates that transmit genetic information from DNA to protein. Accordingly, coding potential has served as the primary criterion distinguishing mRNAs from lncRNAs. Numerous experimental and computational approaches have been developed to enforce this dichotomy. This classification framework implicitly assumes that coding and non-coding functions are mutually exclusive. Once the coding potential is determined, the functional category of a piece of RNA is set.

Recent integrative multi-omic studies have revealed that a fraction of annotated lncRNAs harbor short open reading frames (ORFs) that can be translated to functional micropeptides or microproteins (*45–47*). Over 1000 such micropeptides and microproteins have been identified and validated by mass spectrometry analysis with roles in various cellular processes including mitochondrial homeostasis and myogenesis (*48–52*). These data suggest that transcripts classified as lncRNAs can have coding potentials for short peptides (*46*). In contrast, whether mRNAs themselves exert biologically functions beyond their protein-coding roles remains largely unexplored.

Here, we identify a coding independent function of *Pink1* transcript in skeletal muscle mitochondrial quality control. *Tet2* loss in muscle stem cells (MuSCs) and myofibers reduced *Pink1* transcript abundance, leading to mitochondrial dysfunction and muscle fatigue. Expression of a translation-deficient *Pink1* transcript with all in frame ATG mutated fully rescued the mitochondrial defects in *Tet2* KO cells, suggesting that *Pink1* mRNA functions independently of coding capability and protein production. In wild type (WT) muscle cells, although *Pink1* transcripts was abundant, PINK1 protein was rapidly degraded post translation and undetectable in muscle cells using multiple methods, further supporting the coding and protein independent role of *Pink1* mRNA. Moreover, a portion of *Pink1* mRNA localized to mitochondria and directly interacted with the inner mitochondrial membrane protease YME1L1, facilitating its protease cleavage of OPA1 to maintain mitochondria homeostasis. These findings reveal an intrinsic, protein independent function of a conventional mRNA beyond its traditional protein-coding role. These results challenge the traditional dichotomy between coding and non-coding RNAs and suggest that coding independent functions may be a broader and underappreciated feature of many conventional mRNAs.

## RESULTS

### *Tet2* cKO leads to mitochondria dysfunction in skeletal muscle

We have previously reported that Tet2 regulates the differentiation of MuSCs by reducing DNA methylation at sites proximal to MyoD recognition elements of *Myog* enhancer (*8*). To further investigate the role of Tet2 in skeletal muscle, we generated a MuSC specific *Tet2* conditional knockout (cKO) mouse line *Pax7* CreERT2: *Tet2* flox/flox (Fig. S1A). *Tet2* cKO was induced in 2-month-old mice by Tamoxifen (TAM) injection and muscle function was assessed 4 months later.

*In situ* physiological analysis of skeletal muscle function revealed a decline in muscle force. Both maximal isometric twitch and tetanic forces of Tibialis anterior (TA) muscle were reduced in *Tet2* cKO mice compared with wild type (WT) controls (Fig. 1A). A force-frequency test was performed by stimulating TA muscle at 10-100Hz for 500ms at each stimulus with 3min recovering interval. Whereas WT muscle force plateaued at stimulation frequencies ≥ 80Hz, *Tet2* cKO muscles exhibited reduced force across all frequencies, reaching a plateau at ≥ 40Hz (Fig. 1B). Muscle fatigue was assessed by repetitive stimulation at 30Hz, with force measured at 30sec intervals. WT TA retained approximately 50% of the initial force after 4min of stimulation, whereas *T*et2 cKO muscles reached 50% of the initial force within 2min, indicating markedly reduced fatigue resistance (Fig. 1C). Together, these data demonstrate that Tet2 ablation in MuSCs leads to reduced muscle force and declined fatigue tolerance.

**Fig. 1.**
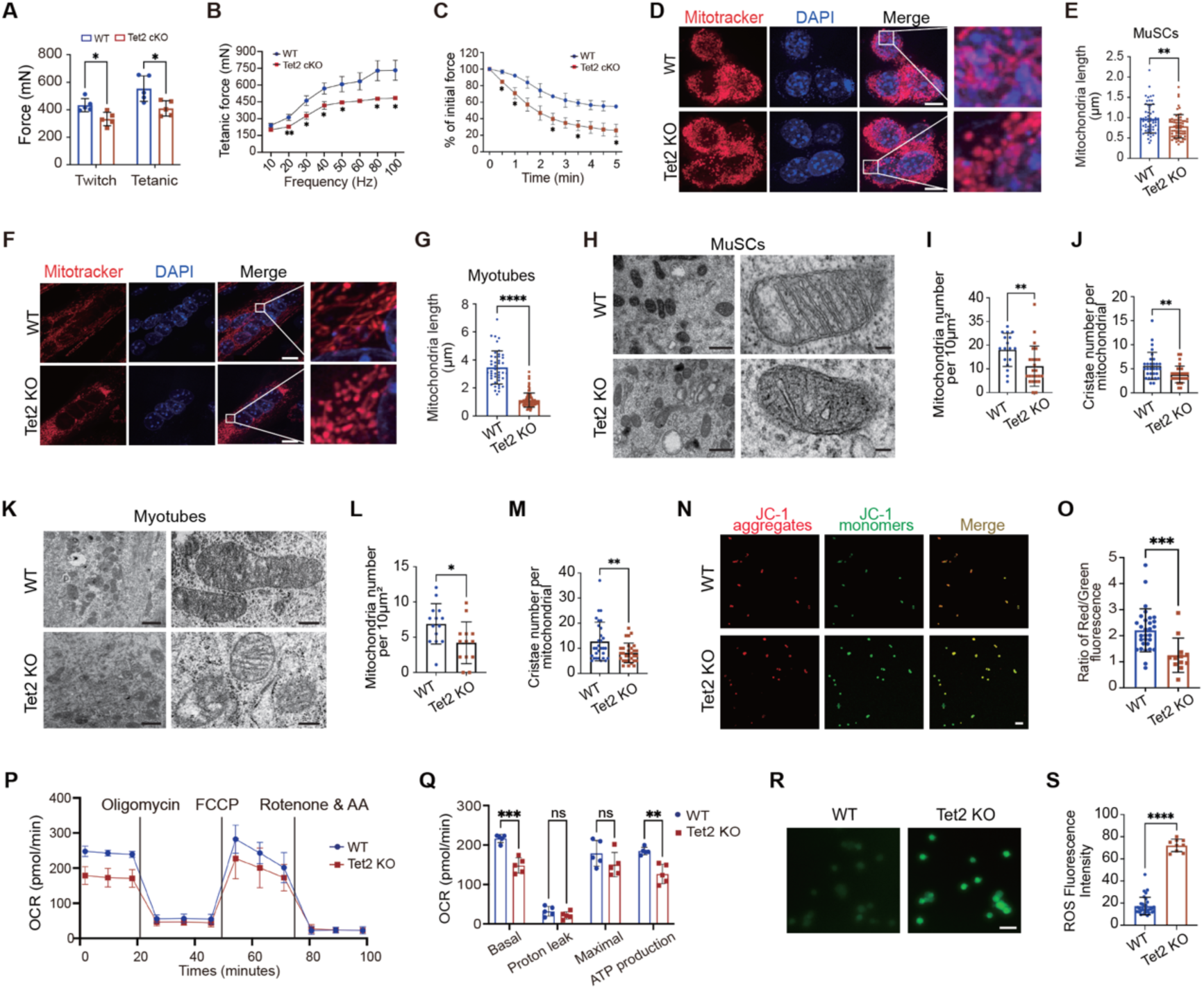
*Tet2* knockout leads to mitochondrial dysfunction in muscle cells. **A.** Quantification of twitch force and tetanic force in WT or *Tet2* cKO TA muscle *in situ* (n=5). **B.** Tetanic Force changes upon frequency gradient stimulations of WT or *Tet2* cKO TA muscle *in situ* (n=5). **C.** Quantification of the fatigue contractions *in situ* (n=5). **D.** Representative images of the mitochondria stained with MitoTracker (red) and DAPI (blue) in WT and *Tet2* KO MuSCs, respectively. Scale bars, 5μm. **E.** Quantification of the mitochondria length in MuSCs. **F.** Representative images of the mitochondria stained with MitoTracker (red) and DAPI (blue) in WT and *Tet2* KO myotubes, respectively. Scale bars, 10μm. **G.** Quantification of the mitochondria length in myotubes. **H.** Representative TEM images of mitochondria in WT and *Tet2* KO MuSCs. Scale bars, 0.5µm and 0.1µm, respectively. **I-J.** Quantification of the mitochondria number (I) and cristae number per mitochondria (J) in WT and *Tet2* KO MuSCs. **K.** Representative TEM images of mitochondria in WT and *Tet2* KO myotubes. Scale bars, 1µm and 0.5µm, respectively. **L-M.** Quantification of the mitochondria number (L) and cristae number per mitochondria (M) in WT and *Tet2* KO myotubes. **N.** Representative images of the mitochondria stained with JC-1 in WT and *Tet2* KO MuSCs, respectively. Scale bars, 50μm. **O.** Quantification of the ratio of red/green fluorescence intensity stained with JC-1 in WT and *Tet2* KO MuSCs. **P.** OCR detected by seahorse assay in WT and *Tet2* KO MuSCs, respectively. **Q.** Quantification of OCR in basal respiration, proton leak, maximal respiration, and ATP production stages (n=5). **R.** Representative images of the mitochondria stained with MitoSox in WT and *Tet2* KO MuSCs, respectively. Scale bars, 50μm. **S.** Quantification of the ROS fluorescence intensity in WT and *Tet2* KO MuSCs. Data are shown as mean ± s.d. Two-tailed unpaired Student’s t-test was used for statistical analysis. ns, not significant. * indicated p < 0.05, ** indicated p < 0.01, *** indicated p < 0.001, **** indicated p < 0.0001.

Mitochondrial dysfunction is a major contributor to muscle fatigue (*53, 54*). Therefore, we examined the morphology of mitochondria from MuSCs and myotubes differentiated from MuSCs using MitoTracker staining. In *Tet2* KO MuSCs, mitochondria appeared fragmented and in shorter length (Fig.1, D to G). To further characterize the mitochondrial defects, the ultrastructure of mitochondria in TA muscle from WT and *Tet2* cKO mice was visualized using transmission electron microscopy (TEM). The mitochondria abundance was reduced by nearly 50% in *Tet2* cKO muscle, and approximately 60% of the remaining mitochondria exhibited swelling morphology, disrupted cristae, and reduced cristae density (Fig. S1, B to E). Consistently, both MuSCs and myotubes from *Tet2* cKO mice displayed reduced mitochondria number and a high prevalence of abnormal mitochondria containing reduced and unstructured cristae (Fig. 1, H to M). Taken together, these results suggest that *Tet2* KO leads to profound mitochondrial structural defects in both MuSCs and myotubes.

We further examined the functions of mitochondria. Mitochondrial membrane potential was evaluated using JC-1 staining, which indicated membrane polarization through a red to green fluorescence ratio. JC-1 forms red aggregates in mitochondria with high membrane potential; whereas it remains in green monomeric form in mitochondria with low membrane potential (*55, 56*). *Tet2* KO MuSCs exhibited a significantly reduced red/green fluorescence ratio compared to WT MuSCs (Fig. 1, N and O), indicating reduced mitochondrial membrane potential. Consistent with the compromised membrane potential, seahorse assay results revealed a marked reduction in oxygen consumption rate (OCR) in *Tet2* KO MuSCs (Fig. 1, P and Q), indicating defective mitochondrial respiration and electron transport chain (ETC) activity. Impaired electron transport is often associated with increased electron leakage and reactive oxygen species (ROS) production (*57, 58*). Indeed, MitoSOX staining demonstrated significantly elevated mitochondrial ROS levels in *Tet2* KO MuSCs (Fig. 1, R and S). Collectively, these results suggest that Tet2 is a critical regulator of mitochondrial integrity and function in muscle cells.

### Tet2 directly activates *Pink1* transcription to maintain mitochondria homeostasis

We next investigated the mechanism by which Tet2 regulates mitochondria homeostasis. Quantification of mitochondrial DNA content revealed no significant differences between WT and *Tet2* KO MuSCs, differentiated myotubes, or TA muscle (Fig. S2A). Consistently, expression levels of mitochondria coded genes were unchanged as shown by RT-qPCR (Fig. S2B). These results suggest that the mitochondrial defects observed in *Tet2* KO muscle cells are unlikely to arise from alterations in the mitochondrial genome, implicating dysregulation of nuclear encoded factors.

Given the pronounced mitochondria fragmentation in *Tet2* KO MuSCs, the expression levels of key regulators of mitochondrial fission and fusion were first examined by RT-qPCR. No significant differences were observed between WT and *Tet2* KO MuSCs (Fig. S2C), suggesting that the defects are not directly caused by transcription alterations in mitochondria biogenesis or dynamics. To identify the potential Tet2 target genes, we further analyzed RNA sequencing and genome-wide methylation sequencing data from *Tet2* KO MuSCs. Total 198 hypermethylated genes were transcriptionally down-regulated in *Tet2* KO MuSCs. Among these, 5 genes encoded mitochondrial localized proteins (Fig. 2A), and were selected for further analysis.

**Fig. 2.**
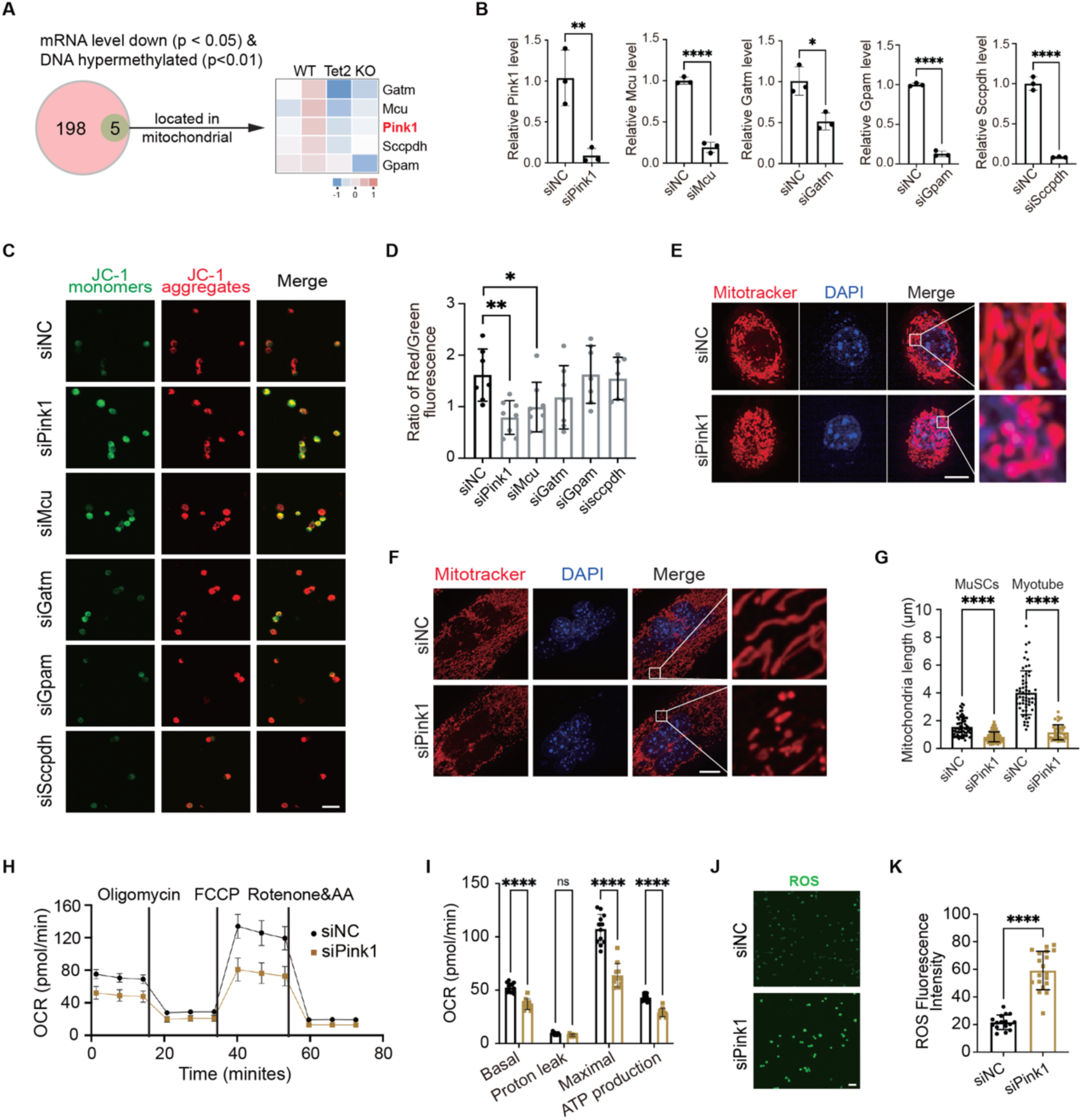
Tet2 regulates mitochondrial homeostasis via epigenetic control of *Pink1* expression. **A.** Venn diagram of the hypermethylated, downregulated genes in *Tet2* KO MuSCs and genes which encode mitochondria localized proteins (left); heatmap diagram of the transcriptional changes of the 5 candidate genes upon *Tet2* KO (right). **B.** RT-qPCR quantification of the mRNA level after scramble (negative control, NC), *Pink1, Mcu, Gatm, Gpam and Sccpdh* siRNA transfection (n=3). **C.** Representative images of the mitochondria stained with JC-1 upon scramble (negative control, NC), *Pink1, Mcu, Gatm, Gpam, and Sccpdh* siRNA transfection in MuSCs. Scale bars, 50μm. **D.** Quantification of the ratio of red/green fluorescence intensity in JC-1 staining. **E.** Representative images of mitochondria stained by MitoTracker (red) and DAPI (blue) in scramble (negative control, NC) and *Pink1* RNAi MuSCs, respectively. Scale bars, 5μm. **F.** Representative images of the mitochondria stained with MitoTracker (red) and DAPI (blue) in scramble (negative control, NC) and *Pink1* RNAi myotubes, respectively. Scale bars, 10μm. **G.** Quantification of the mitochondria length in MuSCs and myotubes. **H.** OCR detected by seahorse assay in scramble (negative control, NC) and *Pink1* RNAi MuSCs, respectively. **I.** Quantification of OCR at basal respiration, proton leak, maximal respiration, and ATP production stages (n=5). **J.** Representative images of the mitochondria stained with MitoSox in scramble (negative control, NC) and *Pink1* RNAi MuSCs, respectively. Scale bars, 50μm. **K.** Quantification of the ROS fluorescence intensity in scramble (negative control, NC) and *Pink1* RNAi MuSCs. Data are shown as mean ± s.d. Two-tailed unpaired Student’s t-test was used for statistical analysis. ns indicated not significant. * indicated p < 0.05, ** indicated p < 0.01, *** indicated p < 0.001, **** indicated p < 0.0001.

Down-regulation of each candidate gene was first validated by RT-qPCR, confirming reduced expression of *Pink1*, *Gatm*, and *Gpam* in *Tet2* KO MuSCs (Fig. S2D). Next, each of the 5 mitochondria located gene was individually knocked down in WT MuSCs by RNAi. These genes were efficiently knocked down as verified by RT-qPCR (Fig. 2B). Assessment of mitochondria membrane potential by JC-1 staining revealed that knockdown of *Pink1*, which encoded a ubiquitin kinase critical for mitophagy and mitochondria quality control (*59*), resulted in the most pronounced reduction of mitochondrial membrane potential (Fig. 2, C and D). We therefore focused subsequent analysis on *Pink1*. Consistent with the phenotype observed in *Tet2* KO MuSCs and myotubes, *Pink1* knockdown induced marked mitochondrial fragmentation (Fig. 2, E to G). Similar to the scenario in *Tet2* KO MuSCs, *Pink1* RNAi led to reduced OCR at basal and ATP production stage of respiration and elevated ROS production (Fig. 2, H to K). These results demonstrate that *Pink1* knockdown phenocopies the mitochondrial structural and functional defects observed in *Tet2* KO muscle cells, suggesting *Pink1* as a key downstream target gene of Tet2.

### Tet2 mediated DNA demethylation activates *Pink1* transcription

To further investigate the mechanism by which Tet2 regulates *Pink1* expression, DNA methylation level at the promoter and enhancer regions of *Pink1* was evaluated using genome wide DNA methylation sequencing data generated previously (*8*). Hypermethylation was detected at 5 enhancer regions of *Pink1* in *Tet2* KO MuSCs and designated E1-E5, respectively (Fig. 3A). The hypermethylation was confirmed by methylation sequencing of each enhancer region. E5 displayed the most pronounced change of hypermethylation in *Tet2* KO MuSCs, increasing from 10.0% to 45.3%, followed by E1, which showed an approximately 40% increase (Fig. 3B). To validate the enhancer identity of these regions, the enrichment of the active enhancer histone mark H3K27Ac and the enhancer associated mark H3K4me1 was evaluated by chromatin immunoprecipitation (ChIP) assays. H3K27Ac was enriched in all 5 enhancers; whereas H3K4me1 enrichment was detected at E1, E3, E4, and E5 (Fig. 3C and D). Based on these profiles, E1, E3, E4, and E5 were selected for further functional analysis. Each region was cloned upstream of the core promoter of *Pink1* to drive the expression of *Luciferase* in a reporter construct and transfected to WT MuSCs (Fig. 3E). Reporter assays revealed that E1, E4, and E5, but not E3, significantly enhanced promoter activity (Fig. 3E, blue bars), suggesting E1, E4, and E5 function as *bona fide* enhancers.

**Fig. 3.**
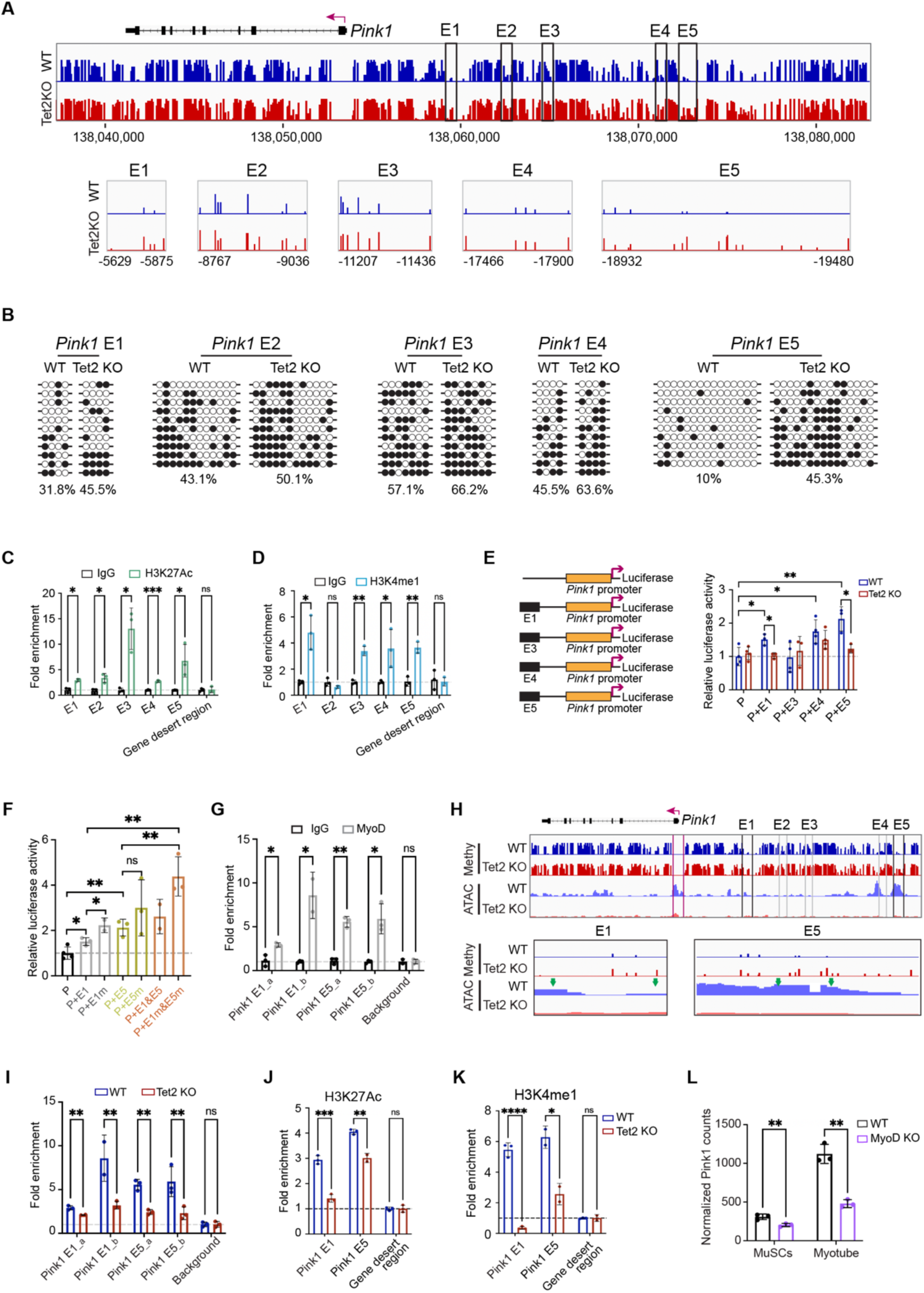
Tet2 mediated DNA demethylation activates *Pink1* transcription. **A.** Methylation profile of the 5’ upstream region of *Pink1* gene in WT or *Tet2* KO MuSCs extracted from genome wide methylation-seq. The top box indicated the methylation profile of *Pink1* gene in WT and *Tet2* KO MuSCs, respectively. The lower box indicated the hypermethylated region in *Tet2* KO MuSCs highlighted in the grey box. And the hypermethylated region was separated into five parts (E1 to E5). **B.** Bisulfite sequencing results of the E1, E2, E3, E4 and E5 enhancer of *Pink1*. White and black circles indicated hypomethylated and hypermethylated CpG sites, respectively. **C-D.** ChIP-qPCR quantification of H3K27ac (C) and H3K4me1 (D) levels in E1-E5 enhancers of *Pink1* (n=3). **E.** Scheme of the luciferase reporter constructs (left) and luciferase activity of the E1, E3, E4 and E5 enhancers (right) (n=3). **F.** Luciferase activity of the E1 and E5 enhancer mutant (n=3). **G.** ChIP-PCR quantification of MyoD binding in *Pink1* E1 and E5 enhancers (n=3). **H.** Methylation and chromatin accessibility profile of the 5’ upstream region of *Pink1* in WT or *Tet2* KO MuSCs extracted from the genome-wide methylation-seq and ATAC-seq. The green arrows indicated the putative MyoD binding sites. **I.** ChIP-qPCR quantification of MyoD recruitment on *Pink1* E1 and E5 enhancer in WT and *Tet2* KO MuSCs (n=3). **J-K.** ChIP-qPCR quantification of H3K27ac (J) and H3K4me1 (K) levels in E1 and E5 enhancers of *Pink1* in *Tet2* KO MuSCs (n=3). **L.** The mRNA level of *Pink1* upon MyoD KO in MuSCs and myotubes. Raw data were from GSA: CRA002490 dataset. Data were shown as mean ± s.d. Two-tailed unpaired Student’s t-test was used for statistical analysis. ns indicated not significant, * indicated p < 0.05, ** indicated p < 0.01, *** indicated p < 0.001, **** indicated p < 0.0001.

To determine whether Tet2 regulates enhancer activity, the Luciferase reporter constructs were transfected to *Tet2* KO MuSCs. Enhancer activity driven by E1 and E5 was markedly reduced upon *Tet2* ablation (Fig. 3E, red bars), suggesting that the activity of E1 and E5 enhancer is Tet2 dependent. To directly test the role of DNA methylation, Cytosines in the CpG islands within E1 and E5 enhancers were mutated to generate methylation resistant enhancers (E1m and E5m). As expected, Luciferase activity driven by the mutant enhancers was higher than that driven by the corresponding WT enhancers (Fig. 3F, grey and green bars), suggesting that DNA hypomethylation relieves transcription repression of *Pink1*. Notably, combining E1 and E5 further enhanced transcription activation, and mutation of CpG sites in both enhancers displayed the strongest transcription activation effect (Fig. 3F, red bars). Together, these results suggest that DNA hypermethylation at E1 and E5 enhancers represses *Pink1* transcription and that Tet2 mediated DNA demethylation is required for efficient transcription activation of *Pink1*.

Sequence analysis of E1 and E5 enhancers revealed 3 putative MyoD recognition elements (Fig. S3). ChIP assays confirmed MyoD occupancy at all predicted sites (Fig. 3G). To assess whether Tet2 dependent DNA methylation influences chromatin accessibility at these regions, ATAC sequencing (ATAC seq) was next performed. *Tet2* KO led to a reduction in chromatin accessibility surrounding MyoD recognition sites within both E1 and E5 enhancers (Fig. 3H). Consistently, MyoD recruitment to these enhancers was significantly diminished in *Tet2* KO MuSCs as determined by ChIP assays (Fig. 3I). Reduced MyoD binding was accompanied by decreased enrichment of the active enhancer histone mark H3K27Ac and H3K4me1 (Fig. 3, J and K), indicating impaired transcription activation. In agreement with the Tet2-MyoD regulatory cascade, *Pink1* expression was down-regulated in *MyoD* KO MuSCs and myotubes (Fig. 3L).

Collectively, these results illustrate an epigenetic mechanism in which Tet2 promotes *Pink1* transcription by demethylating CpG islands within specific enhancer regions, thereby increasing chromatin accessibility and facilitating MyoD recruitment. The Tet2-MyoD axis is essential for robust *Pink1* transcription activation to maintain mitochondria homeostasis in muscle cells.

### Over-expression of *Pink1* mRNA rescues mitochondria defects in *Tet2* KO MuSCs despite absence of PINK1 protein

We next examined whether restoring *Pink1* expression could rescue the mitochondrial defects observed in *Tet2* KO muscle cells. *Tet2* KO MuSCs were infected by lentivirus expressing HA tagged *Pink1*, and mitochondrial function was assessed 4 days post infection (Fig. 4A). As expected, the HA tagged *Pink1* mRNA level increased over 4000-fold after lentivirus infection as indicated by RT-qPCR using primers pairing to HA sequence (Fig. 4B). Concomitantly, mitochondria morphology was markedly improved with increased mitochondria length observed in both *Tet2* KO MuSCs and MuSC differentiated myotubes (Fig. 4, C and D). The defects in mitochondrial membrane potential, bioenergetic capacity, and ROS accumulation were all substantially rescued by *Pink1* overexpression (Fig. 4, E to J). These results suggest that elevating *Pink1* transcript level is sufficient to restore mitochondrial integrity in *Tet2* KO muscle cells, supporting a Tet2-Pink1-mitochondria homeostasis axis.

**Fig. 4.**
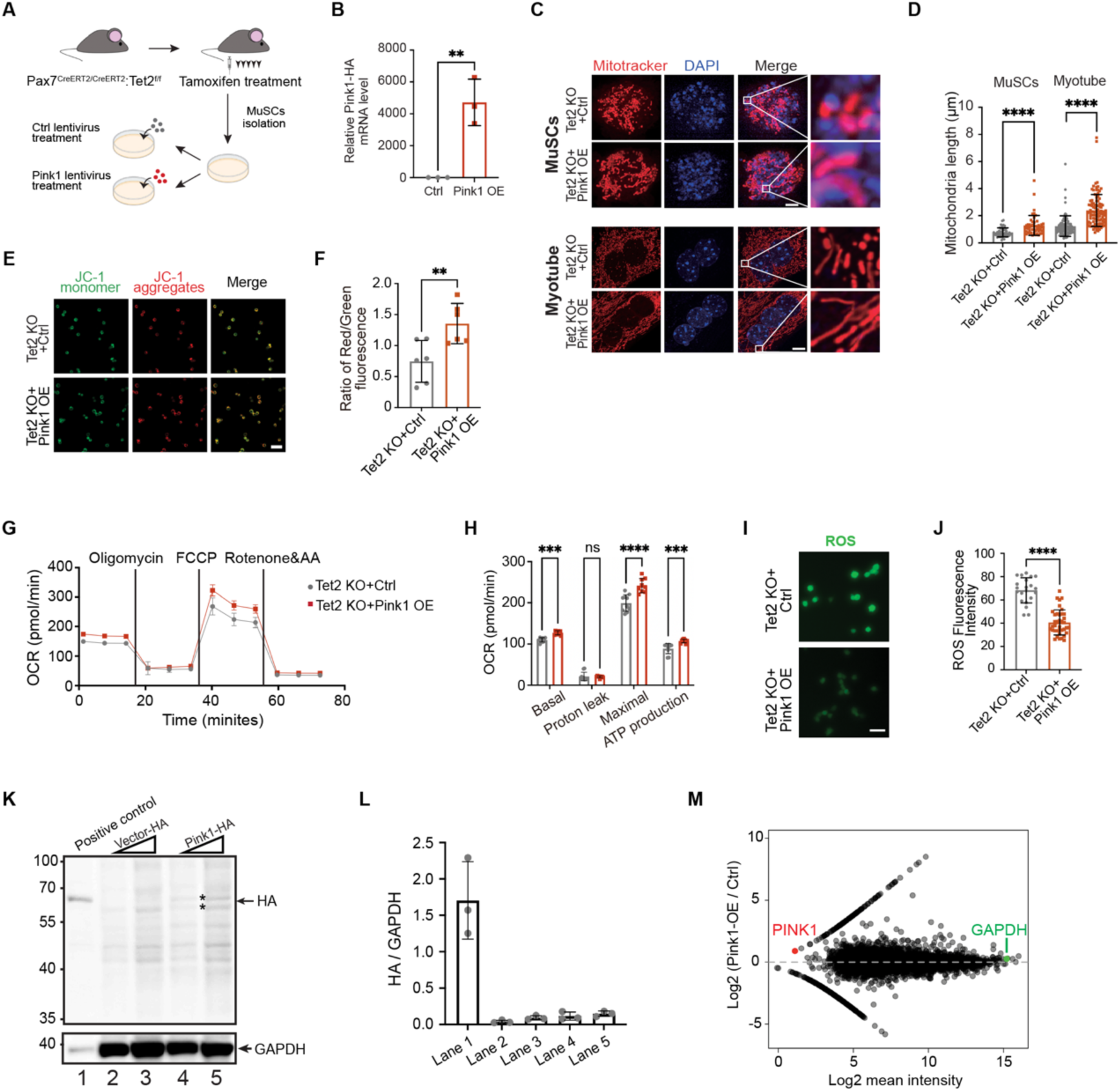
Over-expression of *Pink1* mRNA rescues mitochondria defects in *Tet2* KO MuSCs despite absence of PINK1 protein. **A.** Scheme of the *Pink1* re-expression experiment. **B.** RT-qPCR quantification of the *Pink1-HA* mRNA level (n=3). **C.** Representative images of the mitochondria stained by MitoTracker (red) and DAPI (blue) in control or *Pink1* overexpressed *Tet2* KO MuSCs and myotubes, respectively. Scale bars, 2.5μm and 5μm. **D.** Quantification of the mitochondria length after *Pink1* over-expression in MuSCs and myotubes. **E.** Representative images of mitochondria stained with JC-1 in control or *Pink1* overexpressed *Tet2* KO MuSCs. Scale bars, 50μm. **F.** Quantification of the ratio of red/green fluorescence intensity in JC-1 staining after *Pink1* overexpression. **G.** Seahorse assay showing OCR in control or *Pink1* overexpressed *Tet2* KO MuSCs, respectively. **H.** Quantification of OCR in basal respiration, proton leak, maximal respiration, and ATP production stages (n=10). **I.** Representative images of mitochondria stained with MitoSox in control or *Pink1* overexpressed *Tet2* KO MuSCs, respectively. Scale bars, 50μm. **J.** Quantification of the ROS fluorescence intensity in control or *Pink1* overexpressed *Tet2* KO MuSCs. **K,** Immunoblotting analysis of control and *Pink1-HA* overexpressed *Tet2* KO MuSCs. Two different amount of whole cell extracts were loaded on SDS-PAGE for both control and *Pink1-HA* over-expression samples. HA antibody (CST #3724S) was used. HA tagged PINK1 protein extracted from 293T cells served as a positive control. Non-specific bands were marked by asterisks. **L.** Quantification of Immunoblotting analysis in panel K. **M.** Dot plot displayed the protein intensity difference between control and *Pink1* overexpressed WT MuSCs. Raw data were from mass spectrometry results. Each dot represented a protein, total 8,532 proteins were detected. Data were shown as mean ± s.d. Two-tailed unpaired Student’s t-test was used for statistical analysis. ns indicated not significant, * indicated p < 0.05, ** indicated p < 0.01, *** indicated p < 0.001, **** indicated p < 0.0001.

To further determine the PINK1 protein level after lentivirus infection, immunoblotting was performed using HA antibody. Surprisingly, no specific PINK1 signal was detected using 2 independent HA antibodies (Fig. 4, K and L; Fig. S4, A and B), despite the robust increase in *HA-Pink1* mRNA. To exclude the possibility of tag-specific or HA antibody related artifacts, a second construct encoding FLAG tagged PINK1 was generated. Although mRNA level increased by more than 1800-fold following infection of lentivirus encoding *Flag-Pink1* (Fig. S4C), PINK1 protein remained barely detectable by anti-FLAG immunoblotting (Fig. S4D).

To further rule out the potential limitations of antibody based detection method, we performed quantitative 4D-DIA protein mass spectrometry analysis using whole cell extracts from WT MuSCs over-expressing either empty vector or *Pink1*. A total 8,532 proteins were detected in empty vector or *Pink1* overexpression samples. Notably, PINK1 protein was absent in dataset from empty vector sample, and barely detectable in dataset from *Pink1* overexpression sample (Fig. 4M; Table S1). This finding was unexpected given the relatively high abundance of *Pink1* mRNA in WT MuSCs and thousand-fold increase in Pink1 overexpression MuSCs.

In WT MuSCs, *Pink1* transcript ranked within the top 24.9% of all 19,536 transcripts detected by RNA sequencing (Fig. S4E). By comparison, *Slc6a15*, which encoded the least abundant protein detected by mass spectrometry, exhibited mRNA levels that were only about 1.4% of that of *Pink1* mRNA (Fig. S4E), suggesting proper sensitivity of proteomic analysis. Consistently, proteins encoded by several genes with comparable mRNA level to *Pink1*, including *Nfil3*, *Ptrh2*, *Naa30*, and *Rab18*, were readily detected by the proteomic analysis (Fig. S4F). These results suggest a marked de-coupling between *Pink1* mRNA and PINK1 protein abundance.

A similar disconnect was observed under overexpression conditions. Despite a several thousand-fold increase in *Pink1* mRNA (Fig. 4B), PINK1 protein level remained barely detectable (Fig. 4M). Taken together, these results demonstrate that restoration of mitochondria function in *Tet2* KO MuSCs occurs independently of detectable PINK1 protein, suggesting that *Pink1* mRNA itself rather than its encoded protein mediates the restoration of mitochondrial defects.

### *Pink1* regulates mitochondria homeostasis in a coding independent manner

Previous studies have reported that PINK1 protein can be degraded after translation (*21, 60, 61*). Nuclear and cytoplasmic RNA sequencing data revealed that *Pink1* transcript in MuSCs were properly spliced and efficiently exported to the cytoplasm (Fig. S5A). Polysome sequencing of MuSCs further demonstrated that *Pink1* mRNA associates with actively translating ribosomes, indicating translational engagement (Fig. S5B). We therefore examined whether PINK1 protein was degraded following translation in MuSCs. HA tagged *Pink1* was transfected to WT MuSCs. The transfected cells were subsequently treated with proteasome inhibitor bortezomib (BTZ) for 8hrs prior to immunoblotting analysis. Under these conditions, HA tagged PINK1 protein became readily detected (Fig. S5, C and D). Consistently, quantitative proteomic analysis of whole cell lysates revealed an increase in PINK1 protein abundance following BTZ treatment (Fig. S5E; Table S2). These results suggest that PINK1 translated from *Pink1* mRNA is subject to rapid proteasomal degradation, resulting in negligible steady state protein level in MuSCs.

These observations reveal a paradox in Pink1 function in muscle cells. At the transcript level, *Pink1* is essential for mitochondrial homeostasis, as evidenced by mitochondrial dysfunction following *Tet2* KO or *Pink1* knockdown, and by functional rescue upon *Pink1* overexpression (Fig. 4). In contrast, the encoded PINK1 protein, which is expected to execute the cellular functions of *Pink1* transcript, is largely undetectable under physiological condition, despite pronounced phenotypes upon manipulation of *Pink1* transcript abundance. These seemingly contradictory phenomena suggest that *Pink1* mRNA may have biological functions independent of its protein coding role.

To directly test this hypothesis, a *Pink1* construct, in which all the in frame ATG codons were mutated, was generated, thereby abolishing the protein coding potential while preserving the RNA sequence (Fig. 5A). *Tet*2 KO MuSCs were infected by lentivirus expressing the Flag tagged ATG mutant (Fig. 5B). As expected, no PINK1 protein was detected due to the removal of all in frame ATG codons (Fig. S5, F and G), whereas transcript level of *Pink1* increased by approximately 4000-fold (Fig. 5C). Notably, expression of the translation deficient *Pink1* transcript restored mitochondria morphology and function, as indicated by increased mitochondria length, mitochondrial membrane potential, OCR, and reduced ROS level (Fig. 5, D to K). Taken together, these findings reveal that a translational deficient *Pink1* RNA fully recapitulates the mitochondrial protective effects of the WT *Pink1* mRNA. These data suggest that *Pink1* mRNA itself, rather than the encoded PINK1 protein, is required for maintaining mitochondrial homeostasis in muscle cells, revealing a previously unrecognized coding independent function of a canonical mRNA.

**Fig. 5.**
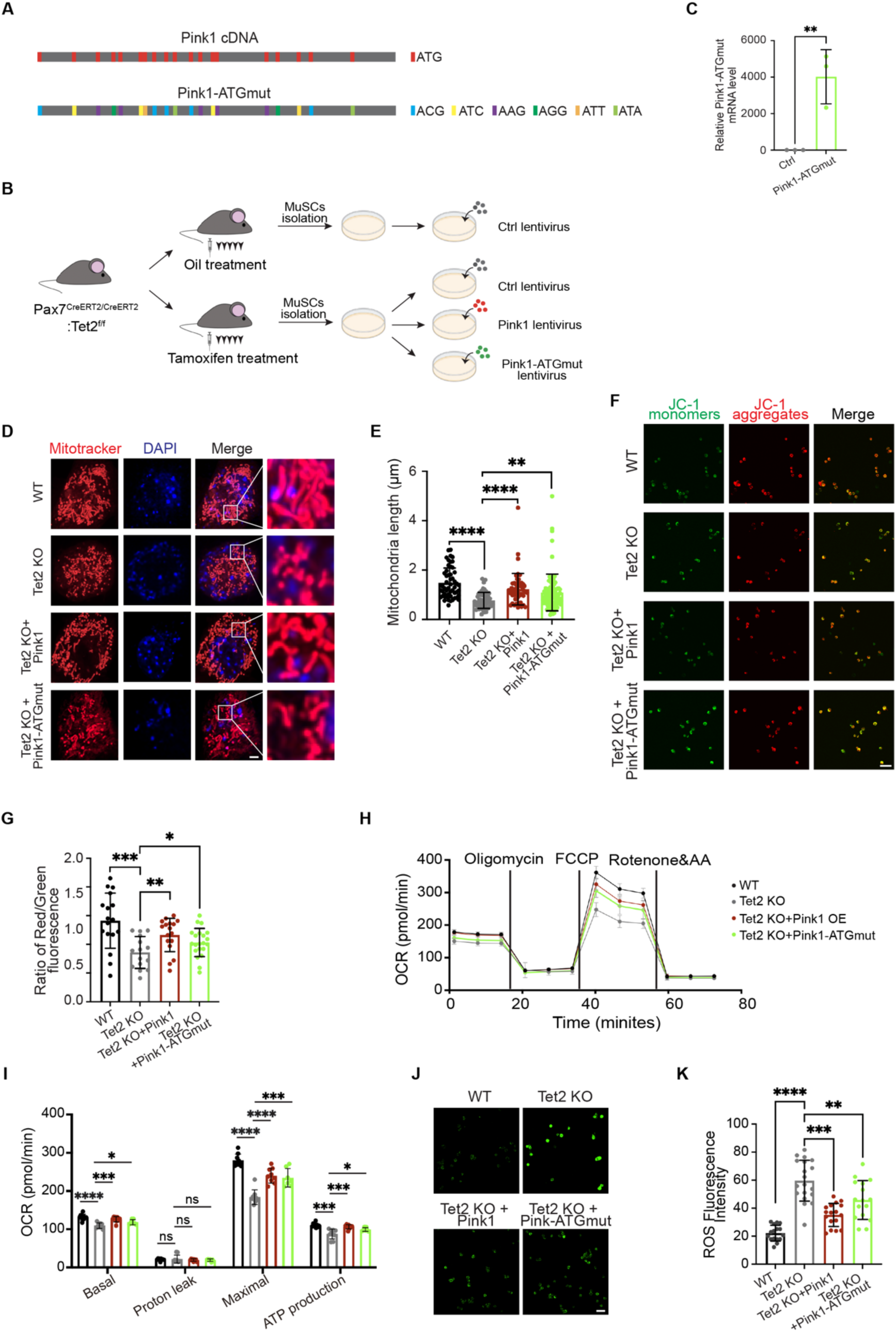
*Pink1* regulates mitochondria homeostasis in a coding independent manner. **A.** Scheme of *Pink1*-*ATGmut* construct. **B.** Scheme of over-expressing *Pink1-ATGmut* in *Tet2* KO MuSCs. **C.** RT-qPCR quantification of the *Pink1-ATGmut* mRNA level (n=3). **D.** Representative images of mitochondria stained by MitoTracker (red) and DAPI (blue) in WT, *Tet2* KO, *Tet2* KO with *Pink1* overexpressed, and *Tet2* KO with *Pink1-ATGmut* overexpressed MuSCs, respectively. Scale bars, 2.5μm. **E.** Quantification of the mitochondria length in WT, *Tet2* KO, *Tet2* KO with *Pink1* overexpressed, and *Tet2* KO with *Pink1-ATGmut* overexpressed MuSCs, respectively. **F.** Representative images of mitochondria stained by JC-1 in WT, *Tet2* KO, *Tet2* KO with *Pink1* overexpressed, and *Tet2* KO with *Pink1-ATGmut* overexpressed MuSCs, respectively. Scale bars, 50μm. **G.** Quantification of the ratio of red/green fluorescent intensity stained by JC-1 in WT, *Tet2* KO, *Tet2* KO with *Pink1* overexpressed, and *Tet2* KO with *Pink1-ATGmut* overexpressed MuSCs, respectively. **H.** Seahorse assay showing OCR in WT, *Tet2* KO, *Tet2* KO with *Pink1* overexpressed, and *Tet2* KO with *Pink1-ATGmut* overexpressed MuSCs, respectively. **I.** Quantification of OCR at basal respiration, proton leak, maximal respiration, and ATP production stages (n=9). **J.** Representative images of mitochondria stained with MitoSox in WT, *Tet2* KO, *Tet2* KO with *Pink1* overexpressed, and *Tet2* KO with *Pink1-ATGmut* overexpressed MuSCs, respectively. Scale bars, 50μm. **K.** Quantification of the ROS fluorescence intensity in WT, *Tet2* KO, *Tet2* KO with *Pink1* overexpressed, and *Tet2* KO with *Pink1-ATGmut* overexpressed MuSCs. Data were shown as mean ± s.d. Two-tailed unpaired Student’s t-test was used for statistical analysis. ns indicated not significant; * indicated p < 0.05, ** indicated p < 0.01, *** indicated p < 0.001, **** indicated p < 0.0001.

### A fraction of *Pink1* mRNA localizes in mitochondria and interacts with YME1L1 to regulate mitochondria homeostasis

We next investigated the mechanism underlying the coding independent function of *Pink1* mRNA. The subcellular localization of *Pink1* mRNA was assessed by RNA fluorescence *in situ* hybridization (RNA FISH) combined with mitochondria staining with mito-dsRed. Surprisingly, approximately 25% of *Pink1* mRNA localized to mitochondria in MuSCs and myotubes (Fig. 6, A and B; Fig. S6, A and B), suggesting a potential direct role in mitochondrial regulation. To identify the proteins that interacted with mitochondria associated *Pink1* mRNA, RNA pull-down assay was performed using biotinylated full length *Pink1* RNA or the translation deficient ATG mutant *Pink1* RNA as baits. Biotinylated antisense *Pink1* RNA (Ctrl-1) and an irrelevant *pTRI-Xef* mRNA (Ctrl-2) served as negative controls. Following incubation with whole cell protein extracts from WT MuSCs, RNA-protein complexes were isolated by streptavidin affinity chromatography and analyzed by mass spectrometry (Fig. 6C). Silver staining showed multiple protein bands specifically enriched in pull-downs using *Pink1* and *Pink1* ATG mutant RNAs as baits, but not in control samples (Fig. 6D). Mass spectrometry analysis identified 326 proteins uniquely interacting with *Pink1* ATG mutant RNA (Fig. 6E). Among the top 20 proteins with high enrichment (average log2 fold change >5), several were mitochondrial proteins (Fig. 6F).

**Fig. 6.**
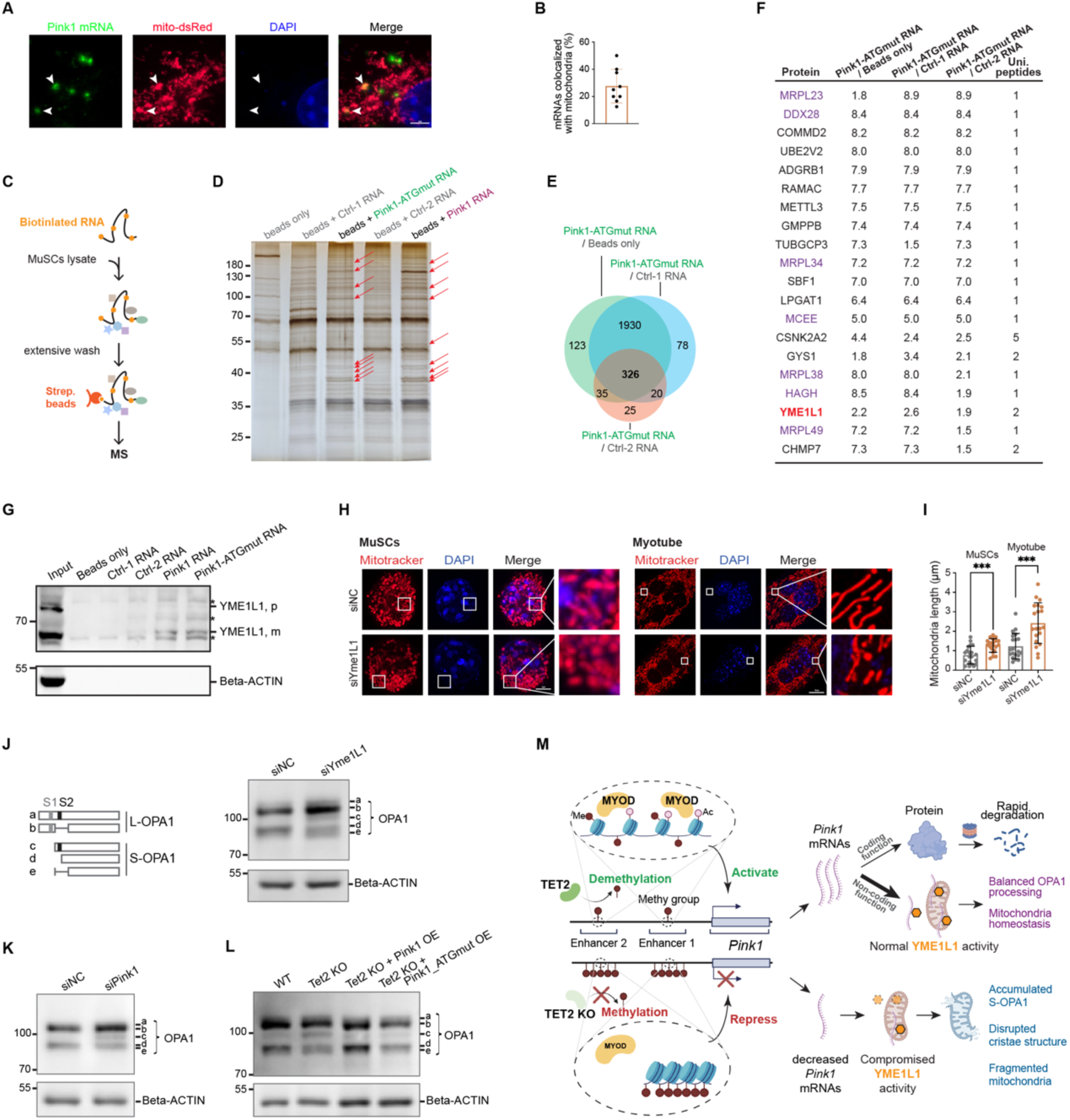
Mitochondria located *Pink1* mRNA interacts with YME1L1 to facilitate OPA1 protein processing. **A.** The location of *Pink1* mRNA in MuSCs assessed by FISH assay. Mitochondria were stained by mito-dsRed. White arrows indicate *Pink1* mRNAs colocalized with mitochondria. Scale bar, 2μm. **B.** Quantification of the percentage of *Pink1* mRNAs localized in mitochondria in MuSCs based on FISH analysis. **C.** Schematic of the experimental workflow of identifying *Pink1*-associated proteins. **D.** Silver staining of proteins pulled down by the *Pink1* mRNA or *Pink1-ATGmut* RNA. Negative controls included blank beads and two unrelated RNA sequences. Red arrows indicated proteins specifically bound by *Pink1* mRNA or *Pink1-ATGmut* RNA. **E.** Ven diagram showing proteins specifically bound by *Pink1-ATGmut* RNA, categorized based on 3 distinct negative controls. Numbers indicated proteins enriched by *Pink1-ATGmut* relative to each control. Overlapping areas highlighted proteins that were enriched by *Pink1-ATGmut* RNA across controls. **F.** Representative proteins specifically enriched by *Pink1-ATGmut* RNA shown in panel E, along with their unique peptides and log2 fold enrichment relative to 3 negative controls. Proteins located in mitochondria were colored in purple. **G.** RNA pulldown followed by Immunoblotting analysis showing the interaction between YME1L1 and *Pink1* mRNA. beta-ACTIN serves as the negative control. YME1L1, p: precursor form of YME1L1. YME1L1, m: mature form of YME1L1. Non-specific bands were marked by asterisk. **H.** Representative images of the mitochondria stained by MitoTracker (red) and DAPI (blue) in scramble (siNC) or *Yme1l1* RNAi MuSCs and myotubes, respectively. Scale bars, 4μm and 10μm, respectively. **I.** Quantification of the mitochondria length in MuSCs and myotubes treated by scramble or *Yme1l1* siRNA, respectively. **J.** Immunoblotting analysis showing OPA1 isoforms in scramble (siNC) and *Yme1l1* RNAi (siYme1L1) MuSCs, respectively. Beta-ACTIN served as an internal control. The schematic diagram on the left illustrated the pattern of intermediate products during proteolytic processing of OPA1. S2 indicated the sites cleaved by YME1L1. L-OPA1: long OPA1 isoforms; S-OPA1: short OPA1 isoforms. *Yme1l1* knockdown led to insufficient YME1L1 level, which impairs OPA1 processing, leading to accumulation of L-OPA1 and the intermediate proteolytic protein product and decreased S-OPA1 d form. **K.** Immunoblotting analysis showing OPA1 isoforms in scramble (siNC) and *Pink1* RNAi (si*Pink1*) MuSCs, respectively. Beta-ACTIN served as an internal control. **L.** Immunoblotting analysis showing OPA1 isoforms in WT, *Tet2* KO, *Tet2* KO with *Pink1* overexpressed, and *Tet2* KO with *Pink1-ATGmut* overexpressed MuSCs, respectively. Beta-ACTIN served as an internal control. **M.** A working model of coding independent function of *Pink1* mRNA maintaining mitochondria homeostasis in skeletal muscle cells.

YME1 like 1 ATPase (YME1L1), a mitochondrial inner membrane protease essential for mitochondria quality control and homeostasis (*62, 63*), emerged as the most prominent mitochondrial interacting protein (unique peptides ≥2) (Fig. 6F). The interaction between YME1L1 and *Pink1* mRNA was validated by RNA pull-down followed by immunoblotting. Both YME1L1 isoforms generated by post-translational cleavage, namely YME1L1p and YME1L1m (*62, 64*), interacted with *Pink1* mRNA, while YME1L1m exhibited higher binding affinity (Fig. 6G). Knockdown of *Yme1l1* by siRNA in both MuSCs and myotubes resulted in shortened and fragmented mitochondria (Fig. S6, C and D; Fig. 6, H and I), phenocopying the defects observed in *Tet2* KO and *Pink1* knockdown muscle cells, suggesting that *Pink1* mRNA-YME1L1 interaction is required for mitochondria homeostasis.

YME1L1 regulates mitochondria inner membrane fusion and cristae organization by proteolytically processing Optic Atrophy 1 (OPA1) protein from its long form (L-OPA1) to short form (S-OPA1) (*63, 65, 66*). Indeed, *Opa1* KO cells displayed fragmented and shortened mitochondria (*67, 68*), similar to the mitochondrial defects observed in *Tet2* KO MuSCs. We next examined whether *Pink1* mRNA interaction affected YME1L1 protease activity. In MuSCs treated by *Yme1l1* siRNA, reduced YME1L1 level led to incomplete OPA1 processing as indicated by the accumulation of multiple intermediate OPA1 species shown in immunoblotting (Fig. 6J; Fig. S6, C and D). In contrast, control MuSCs treated by scramble siRNA showed the expected L-Opa1 and S-Opa1 bands (Fig. 6J). Remarkably, knockdown of *Pink1* mRNA similarly resulted in defective OPA1 cleavage (Fig. 6K), despite intact YME1L1 expression (Fig. S6E). Collectively, these results suggest that *Pink1* mRNA is required for full YME1L1 proteolytic activity.

Consistent with the reduced *Pink1* mRNA level in *Tet2* KO MuSCs, OPA1 processing was likewise impaired in these cells (Fig. 6L), though YME1L1 protein level was not changed (Fig. S6F). Importantly, expression of either WT *Pink1* or the translation deficient ATG mutant *Pink1* RNA fully restored YME1L1 dependent OPA1 cleavage (Fig. 6L), suggesting a coding independent function of *Pink1* transcript. Taken together, these findings suggest that beyond its conventional protein coding role, *Pink1* mRNA localizes to mitochondria and functions as an essential RNA cofactor for YME1L1, enabling proper proteolytic processing of OPA1 in a coding independent manner.

## DISCUSSION

Protein coding and non-coding RNAs have traditionally been classified as functionally distinct entities, with coding capacity viewed as the defining criterion separating mRNAs from non-coding RNAs. Here, by dissecting a Tet2 dependent epigenetic cascade regulating mitochondria functions in muscle cells, we uncovered an unexpected coding independent role for *Pink1* mRNA. *Pink1* transcript, rather than its encoded PINK1 protein, is required to maintain mitochondrial homeostasis in muscle cells. A substantial fraction of *Pink1* mRNA localizes to mitochondria, where it directly interacts with the inner mitochondrial membrane protease YME1L1 to promote efficient OPA1 processing, a key step in regulating mitochondrial inner membrane fusion, cristae organization, and bioenergetic capacity. These findings reveal an intrinsic RNA mediated function of *Pink1* mRNA beyond its canonical role as a protein translation template and blurred the long standing dichotomy between coding and non-coding RNAs (Fig. 6M).

The prevailing view in RNA biology has been that mRNAs and non-coding RNAs represent mutually exclusive functional classes: mRNA act solely through translation, whereas non-coding RNAs exert regulatory functions through RNA based mechanisms. Consequently, coding potential has served as a primary determinant in RNA annotation and functional studies (*23*). Our results challenge this framework by finding that a conventional mRNA can have essential biological functions independently of protein coding capacity and protein production. In this context, *Pink1* mRNA functions as an RNA cofactor of YME1L1 modulating mitochondrial quality control. These findings expand the functional repertoire of mRNAs and broaden our understanding of the centra dogma. Indeed, YME1L1 has previously been reported to have an RNA binding region (RBR) in its ATPase domain (*69*). This RNA binding region might mediate the interaction between Pink1 mRNA and YME1L1.

Importantly, our findings suggest that coding dependent and coding independent functions can coexist within the same transcript. This dual functionality raises the possibility that *Pink1* mRNA is not an isolated case. Indeed, integrative analysis of transcriptomic and proteomic datasets revealed numerous highly abundant mRNAs for which the corresponding protein products were undetectable (Fig. S6G). While some of these discrepancies may reflect technical limitations, others may indicate unrecognized coding independent roles for mRNAs. These observations underscore the need for caution when interpreting results from commonly used techniques to manipulate gene expression such as over-expression, RNAi, and targeted protein degron, which often reflected the combination of RNA mediated and protein mediated effects and cannot be simply interpreted as the functions as the proteins.

## MATERIALS AND METHODS

### Animals

Housing, mating and all experimental protocols for mice used in this study were performed in accordance with the guidelines established by the Institutional Animal Care and Use Committee in Guangzhou National Laboratory. C57BL/6 were obtained from SLRC Laboratory Animal. Tet2 flox/flox mice were kindly provided by Dr. Guoliang Xu (Shanghai Institute of Biochemistry and Cell Biology, CAS, China). Pax7-CreERT2 mice were purchased from Jackson Laboratory (JAX, stock #017763, USA). Tet2 flox/flox mice were crossed with Pax7-CreERT2 mice to generate Pax7-CreERT2: Tet2 flox/flox mice. If not stated differently, 2-month-old male mice were used for all experiments.

Conditional knockout was induced by tamoxifen (Sigma, USA) injection as described previously (*70, 71*). In brief, 10 mg/ml tamoxifen (Sigma, USA) suspended in corn oil was injected intraperitoneally into 2-month-old Pax7-CreERT2: Tet2 flox/flox mice for 5 consecutive days at a dose of 100 mg/kg body weight per day. Littermates of the same genotype were injected with corn oil as a vehicle control.

### Primary MuSCs Isolation, Expansion and Differentiation

Primary MuSCs were isolated as previously described (*8, 72*). Briefly, dissected TA muscles were digested with 10 ml muscle digestion buffer (DMEM containing 1% penicillin/streptomycin (Hyclone, SV30010, USA), 0.125 mg/ml Dispase II (Roche, 04942078001, Germany), and 10 mg/ml Collagenase D (Roche, 11088866001, Germany) for 90 minutes at 37°C. The digestion was stopped by adding 2 ml of FBS (Hyclone, USA). The digested cells were filtered through 70 μm strainers. Red blood cells were lysed by 7 ml RBC lysis buffer (0.802% NH_4_Cl, 0.084% NaHCO_3_, 0.037% EDTA in ddH_2_O, pH 7.2-7.4) for 30s, then filter through 40 μm strainers. After staining with antibody cocktails (AF700-anti-mouse Sca-1 (Thermo Fisher Scientific, 56-5981-82, USA), PerCP/Cy5.5-anti-mouse CD11b (BD Biosciences, 550993, USA), PerCP/Cy5.5-anti-mouse CD31 (BD Biosciences, 562861, USA), PerCP/Cy5.5-anti-mouse CD45 (BD Biosciences, 550994, USA), FITC anti-mouse CD34 (BD Biosciences, 553733, USA), and APC-anti-mIntegrin a7+ (R&D Systems, FAB3518A, USA)), the mononuclear cells were subjected for FACS analysis using Influx (BD Biosciences, USA). The population of PI-CD45-CD11b-CD31-Sca1-CD34+ Integrin a7+ cells was collected. Primary MuSCs were cultured on 0.5 mg/ml Type I collagen (Corning, 354249, USA) coated dishes in proliferation medium (F10 medium containing 15% FBS, 5ng/ml IL-1α (PeproTech, 211-11A, USA), 5ng/ml IL-13 (PeproTech, 210-13, USA), 10ng/ml IFN-γ (PeproTech, 315-05, USA) and 10ng/ml TNF-α (PeproTech, 315-01A, USA), 2.5 ng/ml FGF (R&D Systems, 233-FB-025, USA), and 100ug/ml penicillin/streptomycin (Hyclone, SV30010, USA), and differentiated in differentiation medium (DMEM containing 2% horse serum (Hyclone, SH30074.03, USA) and 100ug/ml penicillin/streptomycin (Hyclone, SV30010, USA)) for 48hr as described previously (*8, 72, 73*).

### *In Vitro* Cell Treatment

MuSCs were cultured and plated at a density of 1.5 ×10^4^ cells/cm^2^ in culture medium. For bortezomib treatment, cells were treated with 1µM Bortezomib (BTZ, MCE, HY-10227, US) for 8hrs, then collected for immunoblotting or mass spectrometry.

### *In Situ* Muscle Force Test

*In situ* hindlimb muscle force analysis was performed with 1300 A 3-in-1 whole animal system (Aurora Scientific, Canada) as previously described (*74, 75*). Mice were anesthetized by 3-bromo-2-fluoro-phenyl methanol (31.2 g/kg body weight) intraperitoneal injection. Mouse TA muscle was exposed by a small incision and half way skin retraction. The distal tendon was carefully severed and sutured to the lever arm of the system, while the ankle was immobilized using clamps. The functional assessment began with three consecutive electrically evoked muscle twitches, followed by a 120sec rest. Subsequently, a series of stimulations at increasing frequencies (10, 20, 30, 40, 50, 60, 80, 100Hz) were applied to induce tetanic contractions. A 180sec rest was set between each frequency. To assess muscle fatigue, the muscles were subjected to a 500ms, 30Hz tetanic stimulation every 2sec over a 4.5min period, during which muscle force was monitored. Electrical stimulations were conducted on the common peroneal nerve in the same leg using subcutaneous platinum electrons with 200μs pulse width. The results were analyzed by DMA software (Aurora scientific, Canada).

### Transmission Electron Microscopy

TA muscle was isolated and immediately fixed in 2.5% glutaraldehyde at 4℃ overnight. Washed once with 0.1M phosphate buffer, then fixed in 1% osmium tetroxide solution at room temperature for 1hr, followed by 2 washes using phosphate buffer. The samples were dehydrated using a series of ethanol gradient (30%, 50%, 70%, 80%, 95%, and 100%) and rinsed twice with acetone. The samples were subsequently embedded in 1:1 mix of acetone: Epon812 resin at room temperature for 2hr, followed by embedding in pure Epon812 resin for 24hr. Thin sections (70nm) were sliced using a Leica EM UC7 ultramicrotome and stained with a solution containing 2% uranyl acetate and lead citrate. Images were acquired using a Gatan US1000 2K CCD camera on FEI Tecnai G2 120V electron microscope.

### Mitochondria Imaging and Function Analysis

Mitochondrial membrane potential and ROS were measured as previously described (*76*). 2×10^4^ MuSCs were seeded on a glass cover slide chamber (Cellvis, China) in proliferation medium or 5×10^4^ cells per well in differentiation medium and allowed to attach for 24hr. For mitochondrial membrane potential staining, living MuSCs or myotubes were incubated with 1μM JC-1 (Abcam, ab113850, UK) for 30min followed by 2 washes with 1×dilution buffer provided with the kit (Abcam, ab113850, UK). For mitochondrial ROS measurement, living cells were incubated with 5μM MitoSOX (Invitrogen, M36008, US) for 30min and washed twice with PBS. After staining, cells were washed with fresh medium 2 times and imaged using Olympus FV3000 microscope (Japan).

For MitoTracker Red staining, live cells were incubated with proliferation or differentiation medium containing 80nM MitoTracker Red (Beyotime, C1035, China) for 30min followed by 3 washes with PBS. Fresh proliferation or differentiation medium were then added. Cells were imaged using the Olympus IXplore SpinSR microscope (Japan). Z-stacks were acquired at a 0.5µm step size covering a total depth of 5µm. And images from 10–20 different fields were captured per imaging dish. Raw z-stack images were deconvolved using the Constrained Iterative plug-in and Advanced Maximum Likelihood algorithm was used for analysis of 5 repeats. Maximum intensity projections were generated from deconvoluted z-stack images. All image analysis were conducted using ImageJ Fiji (1.54 g, National Institutes of Health, US.).

### Seahorse Assay

The Seahorse assay was used to measure OCR (pmol/min)) of MuSCs as previously described (*77, 78*). Briefly, 1.5 × 10^4^ cells/well were plated into collagen type I coated Seahorse XF96 plates (Agilent Technologies, 103793-100, US) and cultured overnight to reach more than 90% confluence. The assay medium was prepared by Seahorse XF DMEM Medium (Agilent Technologies, 103575-100, US) containing 10 mM Glucose, 1mM Pyruvate, and 2mM L-glutamine, pH 7.4. Mitochondrial respiration OCRs were monitored at basal state followed by sequential injection of the 3µM oligomycin, 6µM FCCP, 2.5µM antimycin A and 2µM rotenone to measure OCR at conditions of proton leak, maximal OCR, and ATP production using Agilent Seahorse XF mito-stress kit (Agilent Technologies, 103015-100, US).

### Combined RNA Fluorescence in *Situ* Hybridization (RNA FISH) and Mitochondria Staining

MuSCs were first infected with mito-dsRed lentivirus. 2×10^4^ cells per well were then seeded on a glass cover slide chamber (Cellvis, China) in proliferation medium or 5×10^4^ cells per well in differentiation medium and allowed to attach for 24hr.

The mRNA in situ hybridization was manually carried out by employing PinpoRNA^TM^ multiplex Fluorescent RNA in-situ hybridization kit (GD Pinpoease Biotech *Co.Ltd.*, PIF1000, website: https://www.pinpoease.com, China). A series of short probes (GD Pinpoease Biotech *Co.Ltd.*, 689431-B1, China) sequentially complementary to *Pink1* target RNA sequence covering region 1-1743 bp was designed by patented algorithms (CHINA patent number ZL202110581853.9). Briefly, the cells were fixed by 10% NBF and then the endogenous peroxidase was inhibited by Pre-A solution at room temperature. And then use protease treatment to expose the target RNA molecular and followed by probe hybridization for 2hr at 40℃. Then the signal was amplified sequentially by reaction 1, 2 and 3 while HRP molecular was lastly added at reaction 3 (CHINA patent number ZL202110575231.5). For the last step, a tyramide fluorescent substrate (Akoya Biosciences, Opal^TM^520, US) was added and target RNA was labelled by green fluorescent by TSA assay. Finally, cells were stained with DAPI and imaged using the Olympus IXplore SpinSR microscope (Japan).

### Immunoblotting

Proteins were separated by 10% SDS-PAGE according to standard protocols, transferred to Immobilon-P PVDF Membrane (Millipore, IPVH00005, Germany). After blocking with 5% skim milk in TBST, membranes were incubated with the primary antibodies, YME1L1 (Proteintech, 11510-1-AP, US), OPA1 (HUABIO, HA722673, China), HA (CST, 3724, USA; Abcam, ab1424, UK), Flag (Beijing Ray, RM1002, China), GAPDH (Beijing Ray, RM2002, China), beta-ACTIN (Beijing Ray, RM2001, China) overnight at 4℃. After washing in TBST, Goat anti-mouse (Santa Cruz, SC-2005, US) or anti-rabbit (Santa Cruz, SC-2004, US) IgG-HRP antibodies were applied. The membranes were washed 3 times with TBST and visualized by MiniChemi 580 (Beijing Sage Creation Science Co., Ltd, China).

### Mass Spectrometry

Cells were harvested and washed with PBS twice, followed by lysing in urea extraction buffer (8M urea, 100mM Tris, pH 8.0) containing Protease Inhibitor Cocktail Tablets (Sigma, 5056489001, US). The samples were centrifuged for 20 min at 17,000 g at 4 °C to collect the supernatant. The samples were treated by 10mM dithiothreitol for 30 min at room temperature, 20mM iodoacetamide for 30min at room temperature at dark. Subsequently, the protein mixtures were incubated with Lys-C (enzyme/protein, 1:100, w/w) for 4hr and trypsin (enzyme/protein, 1:50, w/w) for 12hr at 37 °C. The resulting peptide mixtures were quenched with 10% trifluoroacetic acid to a final concentration of 1% and cleaned with home-made C18 StageTips. Peptide samples were separated on nanoElute UHPLC (Bruker Daltonics, Germany) equipped with an Ionopticks Aurora C18 column (250mm x 75μm x 1.6μm) (Ionopticks, Australia), with solvent A (0.1% formic acid in H_2_O) and solvent B (acetonitrile containing 0.1% formic acid) 50°C. Total LC–MS/MS run time per sample was 60min with the following gradient: 0-45 min: 2-22% B; 45-50 min: 22-37% B; 50-55 min: 37-80% B, 55-60 min: 80% B at a flow rate of 300μl/min. The LC was coupled online to a hybrid timsTOF Pro (Bruker Daltonics, Germany) via a CaptiveSpray nano-electrospray ion source. The timsTOF Pro mass spectrometer (Bruker Daltonics, Germany) was operated in positive mode with enabled trapped ion mobility spectrometry (TIMS) at 100% duty cycle (100ms ramp time). Source capillary voltage was set at 1500V and dry gas flow to 3L/min at 180°C. Scan mode was set at data independent parallel accumulation–serial fragmentation (dia-PASEF). The dia-PASEF method cover an m/z range from 350 to 1550, mobility from 0.75 to 1.3 Vs/cm^2^ (1/K0) and 14 dia-PASEF scans, including two IM windows per dia-PASEF scan with variable isolation window widths adjusted to the precursor densities. The collision energy was set to rise linearly over the covered mobility range (for 1/K0 values between 0.6 and 1.6, 20 to 59 eV correspondingly). Total resulting DIA cycle time was estimated to be 1.59 s. The raw data were processed by Spectronaut v19. All data were searched against the mouse proteome database (UniProt, June 2022, 21,992 entries without isoforms). Trypsin was chosen as an enzyme with a maximum of two missed cleavages. Carbamidomethylation on cysteine was considered as a static modification; Oxidation on methionine and protein N-terminal acetylation were selected as dynamic modifications. The other parameters were set as default.

### RNA Interference

WT MuSCs were transfected by siRNAs using Lipofectamine^TM^ 3000 (Invitrogen, L3000015, USA) following the manufacturer’s instructions. Three pieces of siRNA were used for each target gene. The sequences of siRNAs were listed below:

si-*Pink1* sense-1 (5’-3’): GAGGAUUAUCUGAUAGGGCAATT;

si-*Pink1* antisense-1 (5’-3’): UUGCCCUAUCAGAUAAUCCUCTT;

si-*Pink1* sense-2 (5’-3’): CCUGGCUGACUAUCCUGAUAUTT;

si-*Pink1* antisense-2 (5’-3’): AUAUCAAGGAUAGUCAGCCAGGTT;

si-*Pink1* sense-3 (5’-3’): GCUGCAAAUGUGCUGCACUUATT;

si-*Pink1* antisense-3 (5’-3’): UAAGUGCAGCACAUUUGCAGCTT;

si-*Sccpdh* sense-1 (5’-3’): GCUAUUGUAGAGAGCUGAATT;

si-*Sccpdh* antisense-1 (5’-3’): UUCAGCUCUCUACAAUAGCTT;

si-*Sccpdh* sense-2 (5’-3’): GGCGAUAAGGGUAGUUUAATT;

si-*Sccpdh* antisense-2 (5’-3’): UUAAACUACCCUUAUCGCCTT;

si-*Sccpdh* sense-3 (5’-3’): AGUUCAGUAUGCUGCUUAUTT;

si-*Sccpdh* antisense-3 (5’-3’): AUAAGCAGCAUACUGAACUTT;

si-*Gatm* sense-1 (5’-3’): GGUCAAUUAUCAAAGACUATT;

si-*Gatm* antisense-1 (5’-3’): UAGUCUUUGAUAAUUGACCTT;

si-*Gatm* sense-2 (5’-3’): GACUGGUCACUCAAGUAUATT;

si-*Gatm* antisense-2 (5’-3’): UAUACUUGAGUGACCAGUCTT;

si-*Gatm* sense-3 (5’-3’): GCUACAGCUUCCUCCCGAATT;

si-*Gatm* antisense-3 (5’-3’): UUCGGGAGGAAGCUGUAGCTT;

si-*Mcu* sense-1 (5’-3’): GCUUGUCAUUAAUGACUUATT;

si-*Mcu* antisense-1 (5’-3’): UAAGUCAUUAAUGACAAGCTT;

si-*Mcu* sense-2 (5’-3’): GGAAUAUGUUUAUCCAGAATT;

si-*Mcu* antisense-2 (5’-3’): UUCUGGAUAAACAUAUUCCTT;

si-*Mcu* sense-3 (5’-3’): GGAGAAGGUACGAAUUGAATT;

si-*Mcu* antisense-3 (5’-3’): UUCAAUUCGUACCUUCUCCTT;

si-*Gpam* sense-1 (5’-3’): GAAUGAUGUUGCUGAGUAATT;

si-*Gpam* antisense-1 (5’-3’): UUCAUCAGCAACAUCAUUCTT;

si-*Gpam* sense-2 (5’-3’): CAGUGUUACUGAGAAUGUATT;

si-*Gpam* antisense-2 (5’-3’): UACAUUCUCAGUAACACUGTT;

si-*Gpam* sense-3 (5’-3’): GGAACAUUCAGAUUCACAATT;

si-*Gpam* antisense-3 (5’-3’): UUGUGAAUCUGAAUGUUCCTT;

si-*Yme1l1* sense-1 (5’-3’): GAUUAGUAGAAGCACAGAATT;

si-*Yme1l1* antisense-1 (5’-3’): UUCUGUGCUUCUACUAAUCTT;

si-*Yme1l1* sense-2 (5’-3’): AGUACAUUACGUUCCUCUATT;

si-*Yme1l1* antisense-2 (5’-3’): UAGAGGAACGUAAUGUACUTT;

si-*Yme1l1* sense-3 (5’-3’): GGUACAUUAGAUAUGUUCATT;

si-*Yme1l1* antisense-3 (5’-3’): UGAACAUAUCUAAGUGACCTT;

si-Control sense (5’-3’): UUCUCCGAACGUGUCACGUTT;

si-Control antisense (5’-3’): ACGUGACACGUUCGGAGAATT.

### RT-qPCR

Total RNA was extracted by TRIzol Reagent (Thermo, 15596-018, USA), and reverse transcription was performed using Reverse-transcriptase M-MuLV (NEB, M0253L, USA), followed by qPCR analysis using SYRB Green qPCR Mix by QuantStudio6 Flex (Thermo, USA). The expression level of each gene was normalized to that of GAPDH using delta-delta Ct method. Primers used were listed in Table S3.

### Luciferase Reporter Assay

*Pink1* promoter (+84 ∼−282), *Pink1* promoter + E1 (WT or Mutation), *Pink1* promoter + E3, *Pink1* promoter + E4, *Pink1* promoter + E5 (WT or Mutation) and *Pink1* promoter + E1 and E5 (WT or Mutation) were cloned into pGL3 basic vector and transfected to MuSCs using the Lipofectamine^TM^ 3000 (Invitrogen, L3000015, USA), respectively. pGL4.74 containing renilla luciferase was co-transfected serving as internal control. Luciferase activities were measured using Dual-Luciferase Reporter Assay System (Promega, PR-E1910, US) by BioTek Synergy NEO (BioTek, US) following the manufacturer’s instructions 48hr after transfection. Relative luciferase activity was calculated as the ratio of Firefly/Renilla luciferase activity. All experiments were repeated at least 3 times.

### Plasmid Construction

*Pink1* cDNA fused with a HA tag or a Flag tag at the C terminal, mito-dsRed sequence (Addgene, 44386, 4777bp – 5755bp, US) were cloned into the pLenti-CMV-puro (Addgene, 17448, US) lentiviral expression vector using ClonExpress II One Step Cloning kit (Vazyme, #C112, China), respectively. An ATG-mut *Pink1* mutant was generated by site-directed mutagenesis using PrimeSTAR Max DNA Polymerase (Takara, R045B, Japan).

### Conventional Bisulfite Sequencing

Genomic DNA was extracted and treated with EpiArt DNA Methylation Bisulfite Kit (Vazyme, EM101, China) according to the manufacturer’s instructions. Bisulfite-treated DNA was subjected to nested PCR amplification, then purified with the Gel Extraction Kit (TianGen, DP204, China) and cloned into T-Vector pMD20 (Takara, 3270, Japan). Individual clones were sequenced by standard Sanger sequencing. Data were analyzed by BISMA (http://services.ibc.uni-stuttgart.de/BDPC/BISMA/) (*79*). Primers used were listed in Table S4.

### RNA Pull Down

Biotinylated RNAs containing *Pink1*-201 coding sequence or control sequences were synthesized via in vitro transcription using NEB T7 RNA polymerase (Thermo Fisher, AM1344, US) with a biotin labeling mix (Roche, 11685597910, Switzerland), following the manufacturer’s protocol. Approximately 6 × 10^7^ myoblasts were washed with cold PBS and then suspended in 1mL of lysis buffer containing 25 mM Tris-HCl (pH 7.4), 150mM NaCl, 0.2% NP-40, 5% glycerol, 1mM PMSF, 10mM ribonucleoside vanadyl complex (RVC; NEB, S1402S, US), and protease inhibitor cocktail. The suspension was sonicated on ice and centrifuged at 13,000 rpm for 10 minutes and the cell debris pellet was discarded.

The protein-RNA binding reaction mixture was prepared by combining 0.1 parts of 10×RNA-Protein Binding Buffer (0.2M Tris-HCl, pH 7.4, 1.16M NaCl, 20mM MgCl_2_), 0.17 parts of 50% glycerol, 0.23 parts of lysate, and 0.5 parts of H_2_O. To minimize non-specific binding, the mixture was first pre-cleared using streptavidin-coated magnetic beads (Invitrogen, 11205D, US) without RNAs. Subsequently, it was incubated with fresh streptavidin-coated magnetic beads preloaded with biotinylated RNA. Each RNA sample was incubated with 1.5mL of this pre-cleared mixture. The reaction proceeded for 1hr at 4 °C. The beads were then washed five times with wash buffer (50mM Tris-HCl, pH 7.4, 150mM NaCl, 0.05% NP-40) to remove non-specifically bound proteins and cellular debris. Finally, proteins were eluted for subsequent analysis, including silver staining, mass spectrometry, or western blotting.

### ChIP Assay

ChIP assays were performed as previously described (*74, 80*). Briefly, crosslinking was performed in 1% formaldehyde for 10min at room temperature, then stopped by 125mM glycine for 5min. Nuclei were isolated in cell lysis buffer (50mM HEPEs, pH 7.6, 10mM KCl, 1.5mM MgCl_2_, 1mM EDTA, 0.5mM EGTA, 0.5% Triton X-100, 1mM PMSF, 1mM DTT) and chromatin was further extracted from the nuclei using nuclei lysis buffer (50mM HEPEs, pH 7.6, 1mM EDTA, 0.5mM EGTA, 1% Triton X-100, 0.1% deoxycholate, 1mM PMSF, 1mM DTT). The chromatin was sheared to 200–500 bp by sonication using Bioruptor PICO (Diagenode SA, Belgium) and incubated with antibody. Dynabeads^TM^ Protein G (Invitrogen, 10004D, US) were used to capture the precipitated chromatin by antibody. The beads was washed sequentially with sonication buffer (20mM HEPEs, pH 8, 150mM NaCl, 2mM EDTA, pH 8, 0.25% SDS and 1% Triton X-100), high-salt buffer (20mM HEPEs, pH 8, 500mM NaCl, 2mM EDTA, pH 8, 0.1% SDS and 1% Triton X-100), LiCl buffer (10mM HEPEs, pH 8, 250mM LiCl, 1mM EDTA, pH 8, 1% NP-40), TE buffer (1mM EDTA, pH 8, 10mM HEPEs, pH 8) and the crosslink was reversed by incubating in 200mM NaCl, 125μM proteinase K, and 62.5μg/ml RNase A at 65℃ overnight. The DNA fragments were purified by phenol-chloroform extraction and detected by qPCR. The fold enrichment was calculated against IgG ChIP-qPCR. The Primers used were listed in Table S5.

### RNA-seq, WBGS, and ATAC-seq data

All RNA-seq data, WBGS data and ATAC-seq data were published previously with the accession number GSE158649 (*8*).

### Statistical Analysis

The numbers of biological replicates and technical repeats in each experimental group were indicated in figure legends. Error bars represented standard deviation unless noted otherwise. Statistical differences between groups were determined by unpaired two-tailed t-test using GraphPad Prism 8 software (GraphPad Software, USA). Statistical significance was set at p < 0.05. ns indicated no significant difference, * indicated p < 0.05, ** indicated p <0.01, *** indicated p < 0.001, **** indicated p < 0.0001.

## ACKNOWLEDGEMENTS

We thank Dr. Qing Zhang from Proteomics and Metabolomics Core Facility, Guangzhou National Laboratory for helps with the sample preparation, LC–MS/MS experiment and data analysis; cell biology facility of Shanghai Institute of Biochemistry and Cell Biology and Guangzhou Laboratory for helps with MuSCs sorting; molecular biology facility of Shanghai Institute of Biochemistry and Cell Biology for helps on transmission electron microscopy analysis; molecular biology facility of Guangzhou National Laboratory for helps on seahorse analysis.

## Author contributions

Conceptualization, P.H.; Investigation, X.-F.W., Y.-Y.L., R.-M. M., and H.-Y.W.; Data analysis, X.-F.W., P. H., H. C., W.-L. Y., X.-J. L., and G.-L.Xu.; Writing – Original Draft, X.-F.W. and P.H.; Writing – Review & Editing, P.H. and X.-F.W.; Supervision, P.H.; Funding Acquisition, P.H.

## Data and materials availability

All RNA-seq data, WBGS data and ATAC-seq data were published previously with the accession number GSE158649 (*8*). All data generated in this study are available in the main text or the supplementary materials. This paper does not report original code. Addgene catalog numbers for plasmids used in this study are noted in the methods. All materials used in this study will be made available upon request.

---

**Fig. S1.**
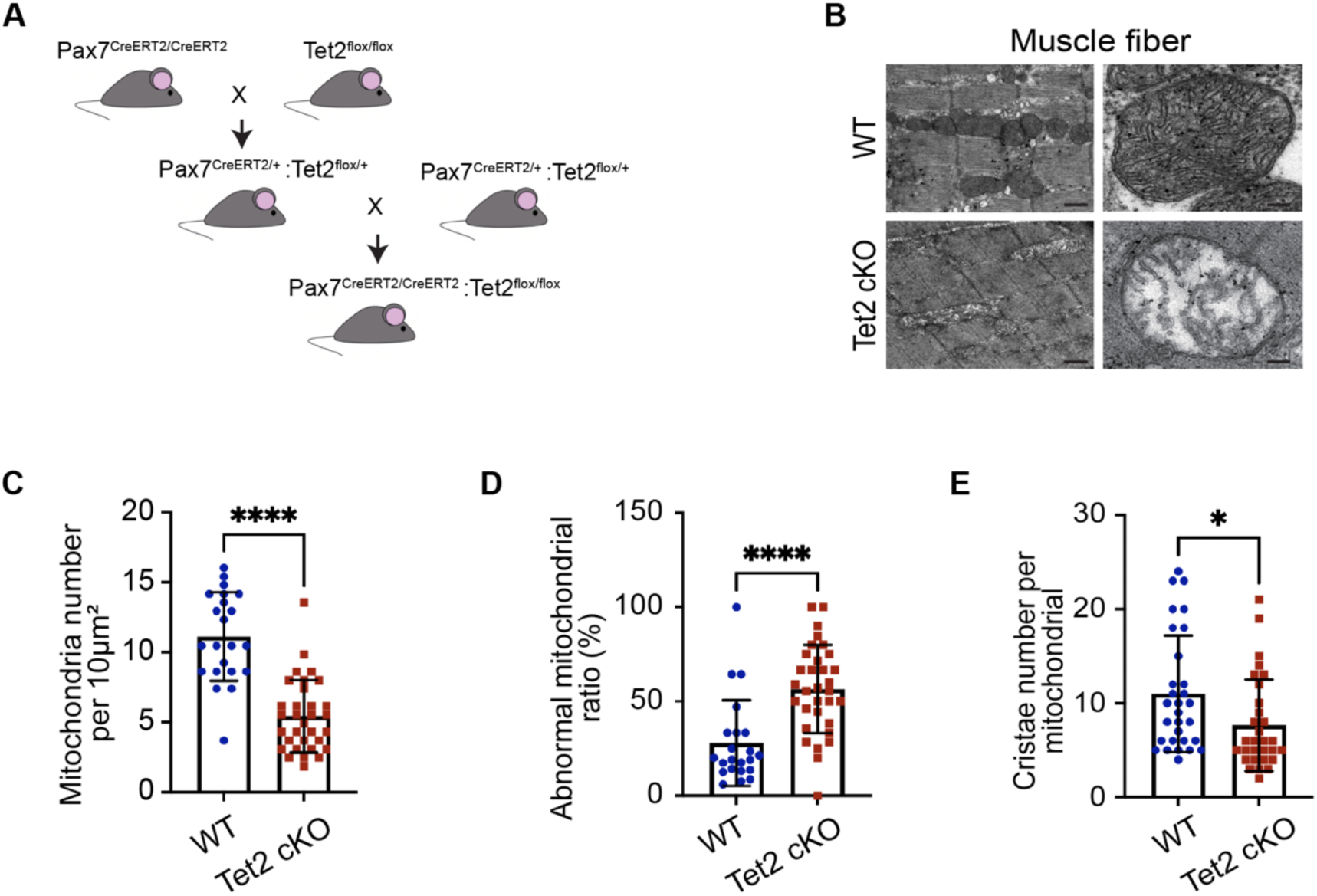
MuSC specific *Tet2* cKO display obvious muscle mitochondria morphology changes 4-months after KO. Related to Fig. 1. **A.** Mating scheme to generate Pax7 CreERT2: Tet2 flox/flox mice. **B.** Representative TEM images of intermyofibrillar mitochondria in WT and *Tet2* cKO TA muscle. Scale bars, 0.5µm and 0.1µm, respectively. **C-E,** Quantification of the mitochondria number (C), abnormal mitochondria ratio (D) and cristae number per mitochondria (E) in TA muscle. Data were shown as mean ± s.d. Two-tailed unpaired Student’s t-test was used for statistical analysis. ns indicated not significant, * indicated p < 0.05, ** indicated p < 0.01, *** indicated p < 0.001, **** indicated p < 0.0001.

**Fig. S2.**
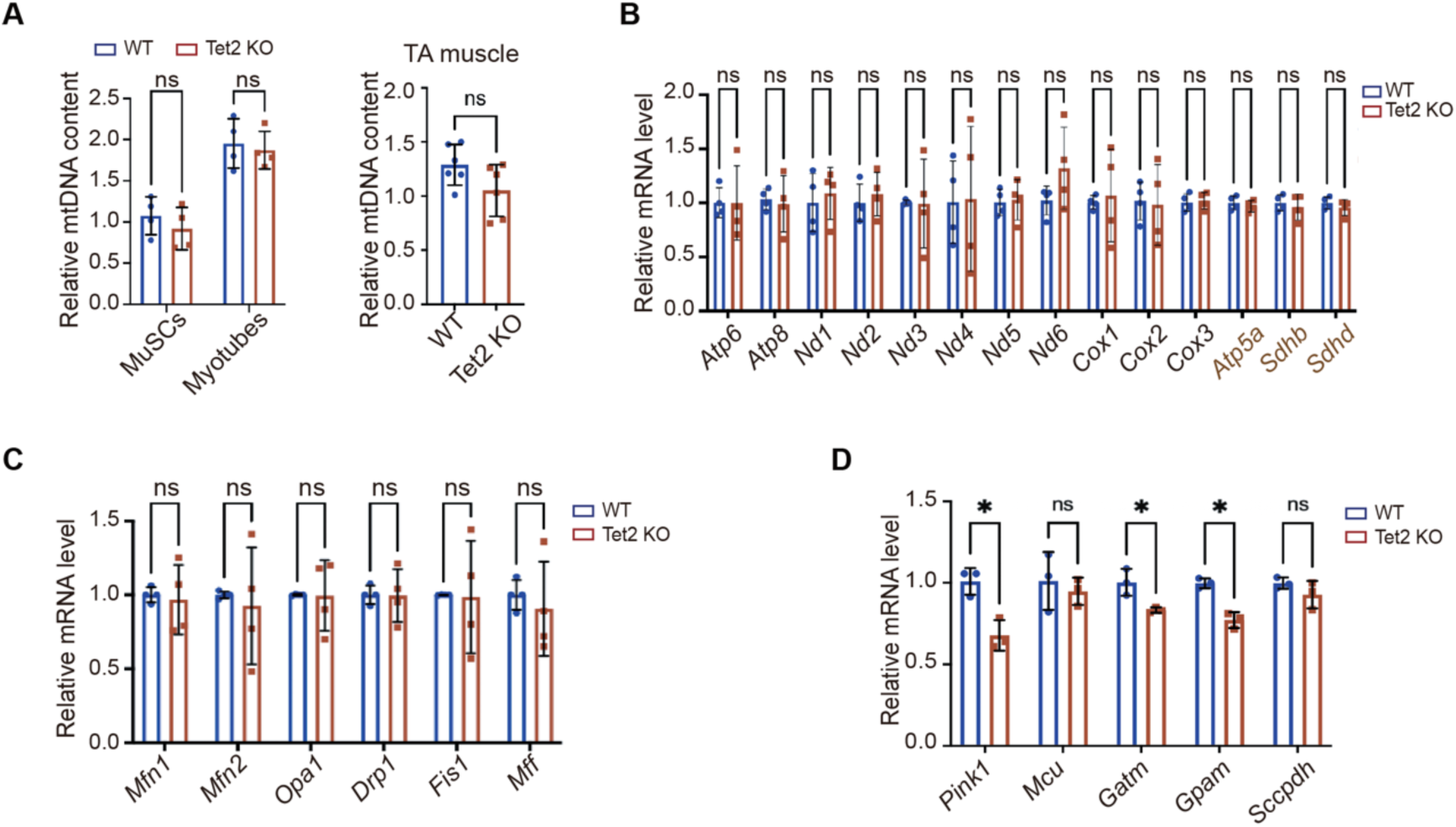
The expression of mitochondria coded genes were not affected by *Tet2* KO. Related to Fig. 2. **A.** Quantification of mtDNA content in MuSCs, myotubes, and TA muscles, respectively. **B.** RT-qPCR quantification of the mitochondrial-encoded (*Nd1-6*, *Cox1-3*, *Atp6*, and *Atp8*) and nuclear-encoded (*Atp5a*, *Sdhb*, *Sdhd*) mRNAs (n=4). **C.** RT-qPCR quantification of the mRNAs encoding key mitochondrial fusion (*Mfn1*, *Mfn2*, *Opa1*) and fission (*Drp1*, *Fis1*, *Mff*) proteins (n=4). **D.** RT-qPCR quantification of 5 candidate mRNAs (n=3). Data were shown as mean ± s.d. Two-tailed unpaired Student’s t-test was used for statistical analysis. ns indicated not significant, * indicated p < 0.05.

**Fig. S3.**
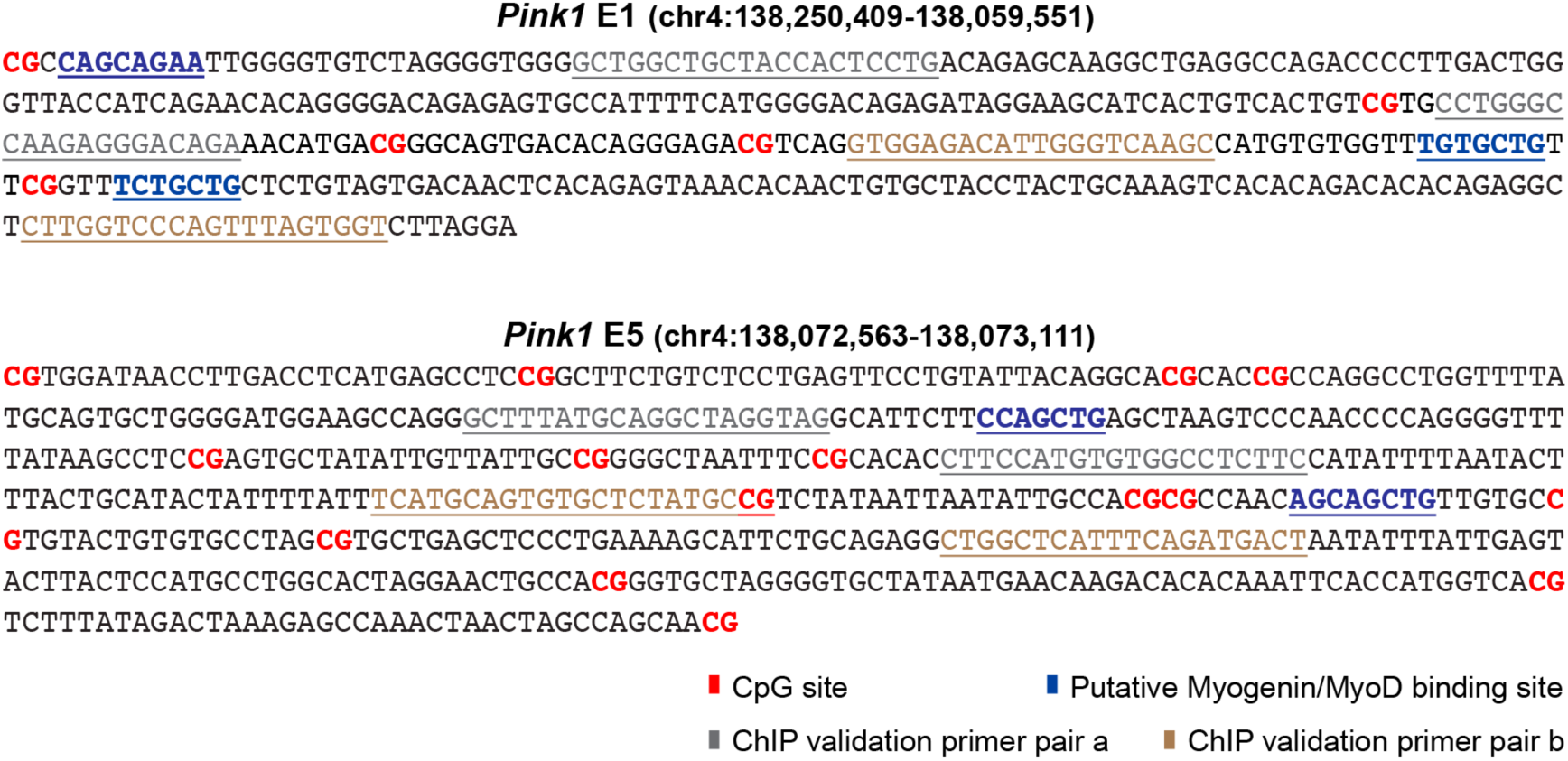
DNA sequence of *Pink1* E1 and E5 enhancers. Related to Fig. 3. CpG sites, putative MyoD binding sites, and the positions of primers used for ChIP-qPCR were labelled by different colors.

**Fig. S4.**
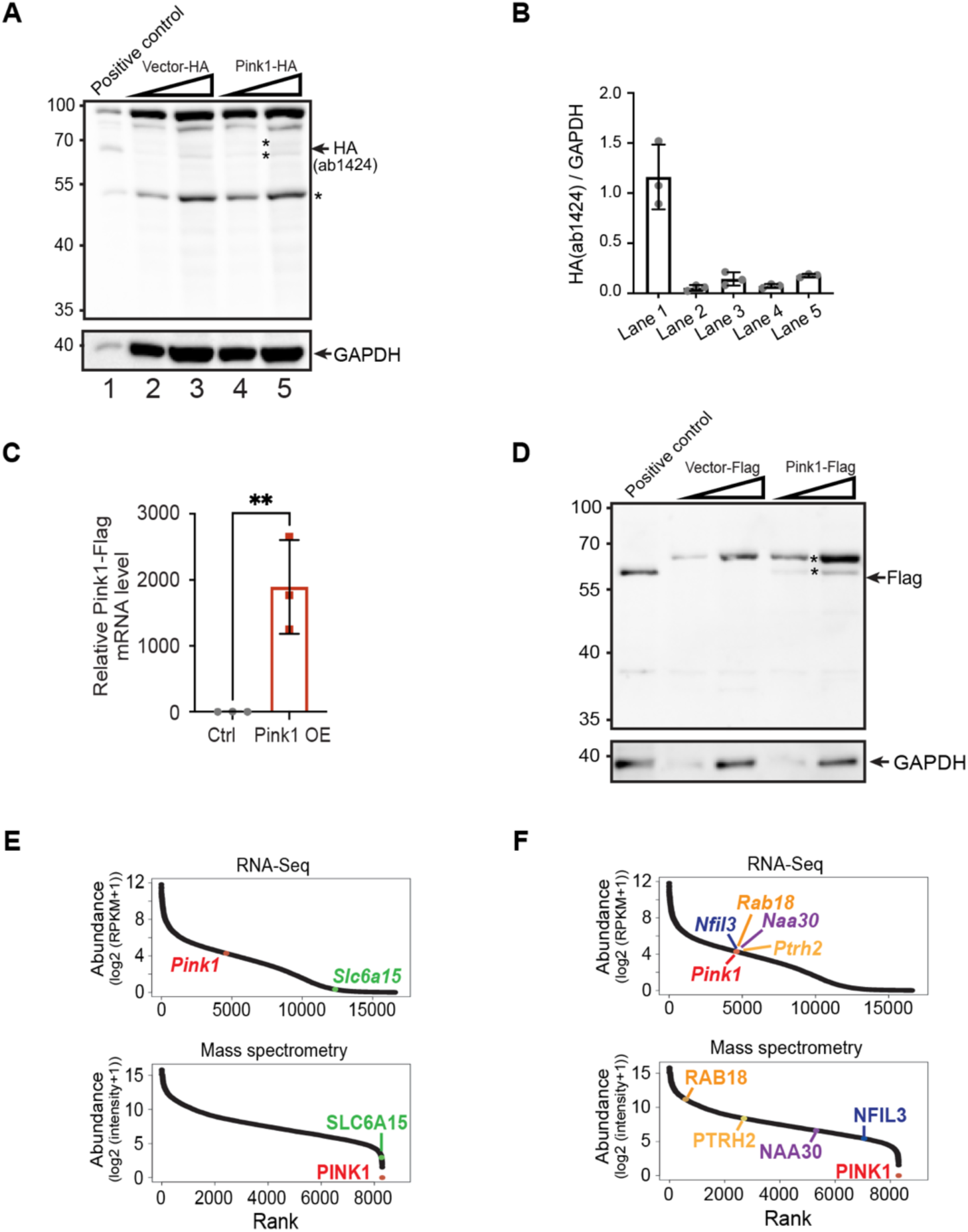
PINK1 protein was undetectable in MuSCs. Related to Fig. 4. **A,** Immunoblotting analysis of control and *Pink1-HA* overexpressed *Tet2* KO MuSCs. Two different amount of whole cell extracts were loaded on SDS-PAGE for both control and *Pink1-HA* over-expression samples. HA antibody (ab1424) was used. HA tagged PINK1 protein extracted from 293T cells served as a positive control. The non-specific bands were marked by asterisks. **B,** Quantification of Immunoblotting analysis in panel A. **C.** RT-qPCR quantification of the *Pink1-Flag* mRNA level (n=3). **D.** Immunoblotting analysis of control and *Pink1-FLag* over-expressed *Tet2* KO MuSCs. Two different amount of whole cell extracts were loaded on SDS-PAGE for both control and *Pink1-FLag* over-expression. Flag antibody was used. Flag tagged PINK1 protein extracted from 293T cells served as a positive control. The non-specific bands were marked by asterisks. **E,** Cumulative rank plot comparing the mRNA abundance of *Pink1*, *Slc6a15* in RNA-seq dataset and protein abundance of PINK1, SLC6A15 in mass spectrometry dataset. **F.** Cumulative rank plot comparing the mRNA abundance of *Pink1*, *Nfil3, Rab18, Naa30, Ptrh2* in RNA-seq dataset and protein abundance of PINK1, NFIL3, RAB18, NAA30, PTRH2 in mass spectrometry dataset.

**Fig. S5.**
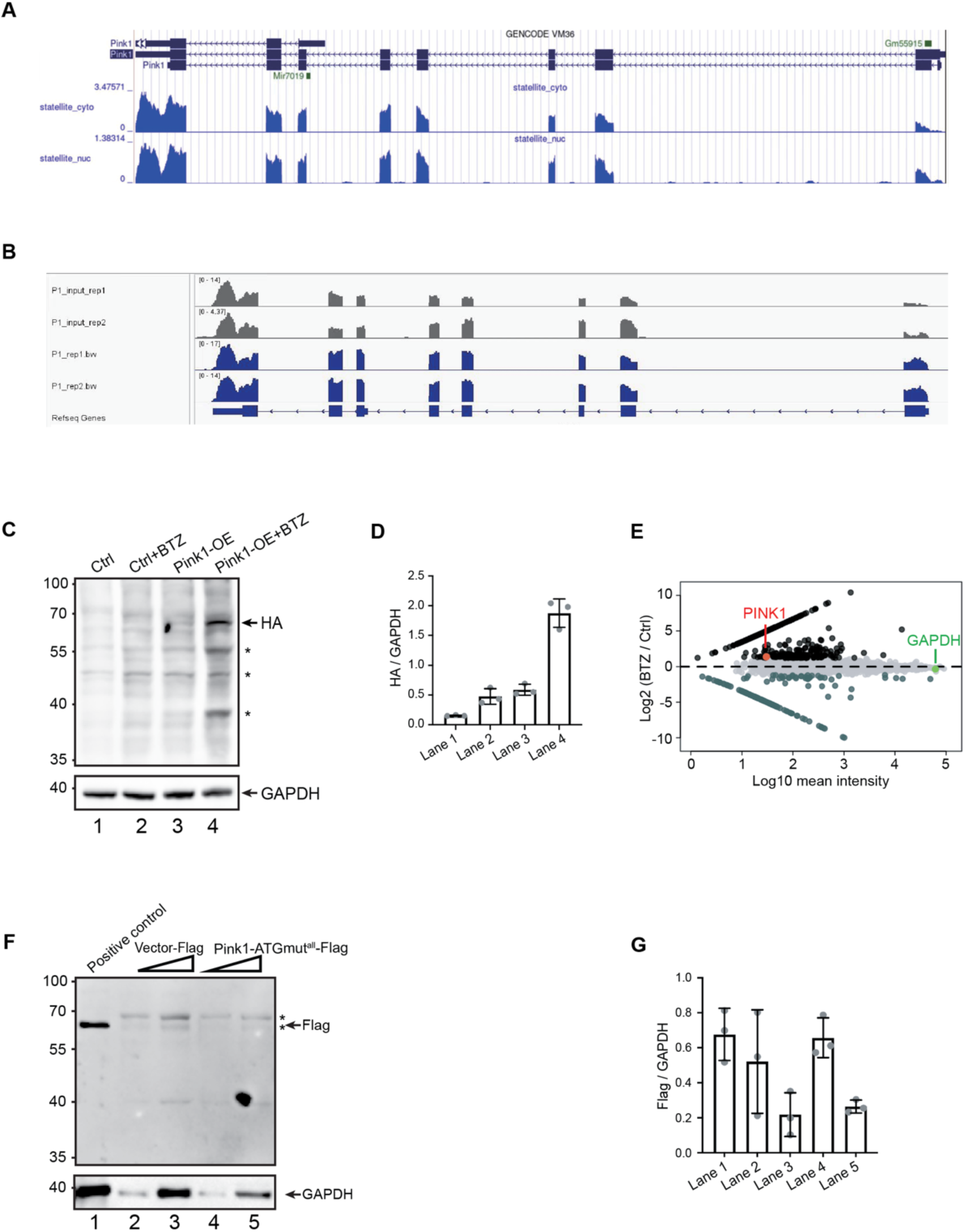
PINK1 protein underwent rapid degradation after translation. Related to Fig. 5. **A.** Fractionated RNA-seq profiles showing the nuclear (nuc) and cytoplasmic (cyto) distribution of *Pink1* mRNA in WT MuSCs. **B.** IGV screenshots showing that *Pink1* mRNAs were enriched in polysome fractions in WT myoblast samples, based on two replicates. **C.** Immunoblotting of PINK1 protein in control or *Pink1-HA* overexpression MuSCs with or without BTZ treatment using HA antibody (CST #3724S). The non-specific bands were marked by asterisks. **D.** Quantification of Immunoblotting analysis in panel C. **E.** Dot plot displayed the protein intensity difference between control and BTZ treated WT MuSCs. Raw data were from mass spectrometry results. Each dot represented a protein, total 9,504 proteins were detected. Black-colored dots: proteins with log2 (BTZ/Ctrl) > 1.2. Darkgreen-colored dots: log2 (BTZ/Ctrl) < −1.2. **F.** Immunoblotting analysis of control and *Pink1-ATGmut-Flag* over-expressed *Tet2* KO MuSCs. Two different amounts of whole cell extracts were loaded on SDS-PAGE. Flag antibody was used. Flag tagged PINK1 protein extracted from 293T cells served as a positive control. The non-specific bands were marked by asterisks. **G.** Quantification of Immunoblotting analysis in panel A. Data were shown as mean ± s.d.

**Fig. S6.**
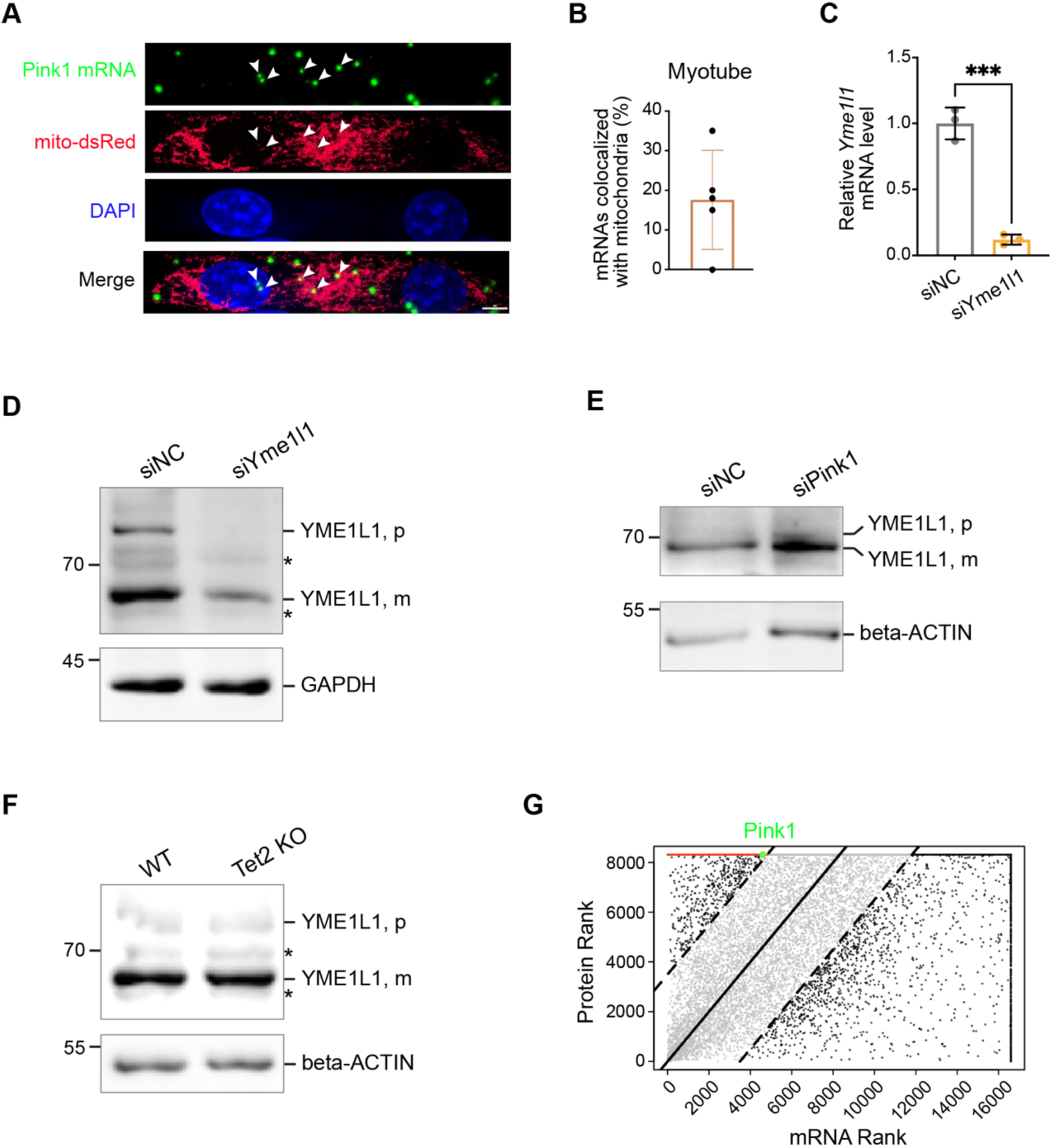
*Pink1* mRNA localized in mitochondria. Related to Fig. 6. **A.** The location of *Pink1* mRNA in myotubes assessed by FISH assay. Mitochondria were marked by mito-dsRed. White arrows indicate *Pink1* mRNAs colocalized with mitochondria. Scale bar, 4μm. **B.** Quantification of the percentage of *Pink1* mRNAs localized in mitochondria in myotubes based on FISH assay. **C.** RT-qPCR quantification of the mRNA level after scramble (negative control, NC), *and Yme1l1* siRNA knockdown (n=3). **D.** Immunoblotting analysis showing precursor (p) or mature (m) form of YME1L1 in scramble (siNC) and *Yme1l1* RNAi (si*Yme1l1*) MuSCs, respectively. GAPDH served as an internal control. **E.** Immunoblotting analysis showing precursor (p) or mature (m) form of YME1L1 in scramble (siNC) and *Pink1* RNAi (si*Pink1*) MuSCs, respectively. Beta-ACTIN served as an internal control. **F.** Immunoblotting analysis showing precursor (p) or mature (m) form of YME1L1 in WT and *Tet2* KO MuSCs, respectively. Beta-ACTIN served as an internal control. **G.** Dot plot illustrating mRNA abundance rank (the mRNA with the highest abundance ranks 1, the mRNA with the lowest abundance ranks 16,590; mRNAs with same RPM values have the same rank) and protein abundance rank (the protein with the highest abundance ranks 1, the protein with the lowest abundance ranks 8,300, proteins with same abundance have the same rank) for 17,135 protein-coding genes detected either by RNA-seq or spectrometry in WT MuSCs. Black-colored dots indicated genes with discrepant mRNA level and protein level. Red-colored dots indicated genes with high mRNA level but undetectable protein level. Pink1 is highlighted with a green dot.

