## Supplementary material for "The non-coding facet of *Pink1* mRNA regulates mitochondria homeostasis": Table S3 to S5

Science
AAAS

### Supplementary Materials for

#### The non-coding facet of *Pink1* mRNA regulates mitochondrial homeostasis

Xiaofen Wu *et al.*

**The PDF file includes:**

Tables S3 to S5

**Other Supplementary Materials for this manuscript include the following:**

Tables S1 to S2

#### Table S3. Primer sequences for RT-qPCR.

| Gene | Forward (5'-3') | Reverse (5'-3') |
|---|---|---|
| Tet2 | CAATAATCAGGTTGAATTTGAAC | GAGGGTGACCACCACTGTACTGC |
| Pink1 | CGACAACATCCTTGTGGAGTGG | CATTGCCACCACGCTCTACACT |
| Gatm | TGGTGCCAAGTGGACAACAGCA | GCAAGGCTCAAACTCAGTCGTC |
| Mcu | GAGCCGCATATTGCAGTACGGT | AAACACGCCGACTGAGTCAGAG |
| Sccpdh | GCACTCAAGTGAAGGGACCAGA | GGAGAAAGCTGCTCCAGGTGTA |
| Gpam | GCAAGCACTGTTACCAGCGATC | TGCAATCAGCCTTCGTCGGAAG |
| Yme1l1 | GCCAGATGTGAAGGGTCGAACT | ACTCTGCTCCAGAGAACCCAAC |
| Mfn1 | CCAGGTACAGATGTCACCACAG | TTGGAGAGCCGCTCATTCACCT |
| Mfn2 | GTGGAATACGCCAGTGAGAAGC | CAACTTGCTGGCACAGATGAGC |
| Opa1 | TCTCAGCCTTGCTGTGTCAGAC | TTCCGTCTCTAGGTTAAAGCGCG |
| Drp1 | GCGAACCTTAGAATCTGTGGACC | CAGGCACAAATAAAGCAGGACGG |
| Fis1 | GCTGGTTCTGTGTCCAAGAGCA | GACATAGTCCCGCTGTTCCTCT |
| Mff | CTACTCGTAGGGCTTACCAGCA | GCTCCTTCAATGGCTGCATCTAG |
| Nd1 | CAGCCGGCCCATTCGCGTTA | AGCGGAAGCGTGGATAGGATGC |
| Nd2 | TCCTCCTGGCCATCGTACTCAACT | AGAAGTGGAATGGGGCGAGGC |
| Nd3 | ACCCTACAAGCTCTGCACGCC | GCTCATGGTAGTGGAAGTAGAAGGGCA |
| Nd4 | TCGCCTACTCCTCAGTTAGCCACA | TGATGATGTGAGGCCATGTGCGA |
| Nd5 | TCGGAAGCCTCGCCCTCACA | AGTAGGGCTCAGGCGTTGGTGT |
| Nd6 | AATACCCGCAAACAAAGATCACCCAG | TGTTGGGGTTATGTTAGAGGGAGGGA |
| Cox1 | GCACTGGTGGATGCCTTCT | TCTCTCGGGACTCCTTGATGA |
| Cox2 | AGTTGATAACCGAGTCGTTCTGCCA | TCGGCCTGGGATGGCATCAGT |
| Cox3 | ACCTACCAAGGCCACCACACTCC | GCAGCCTCCTAGATCATGTGTTGGT |
| Atp6 | GCTCACTCGCCCACTTCCTTCC | GCCGGACTGCTAATGCCATTGGTT |
| Atp8 | ATGCCACAACTAGATACATCAACA | GGGGTAATGAATGAGGCAAA |
| Atp5a | CTTGACCTTCCTTTGCGCTC | GCACCAACAAAGGATGACCC |

| | | |
|---|---|---|
| Sdhb | TGACGTCAGGAGCCAAAATGG | CCTGAAACTGCAGGCCGACT |
| Sdhd | GGTTGTCAGTGTTCTGCTCTTGG | GTCGGTAACCACTTGTCCAAGG |
| Gapdh | GCCAGCCTCGTCCCGTAGACA | CAACAATCTCCACTTTGCCACTGC |

**Table S4. Primer sequences for conventional bisulfite sequencing.**

| Name | Forward (5'-3') | Reverse (5'-3') |
|---|---|---|
| Pink1-E1-1 | GGTTATTATTAGAATATAGGGGATAGAGAGTG | ACACAATTATATTTACTCTATAAATTATCACTAC |
| Pink1-E1-2 | AGTGTTATTTTTATGGGGATAGAGATAGGAAG | CTATAAATTATCACTACAAAACAACAAAAACC |
| Pink1-E2-1 | TGTTTAGTTATTGTTTTGAATATGAATAGTGG | CCTAAAACTCAAACCCAAATTCTAATAAAAAT |
| Pink1-E2-2 | TATTTTTATTAGAATTTGGGTTTGAGTTTTAG | ATATCCCCTAAAAAACACTATAATTATTAACC |
| Pink1-E2-3 | TTTTTGAGTATTAGGGGATTGAAGGAGGTAG | CAAACAAAAACAAACATTACTCAACAATAACTTC |
| Pink1-E3 | TTTTGTTGTATTTATTGATAAAGGG | AACCAACTATAAACTTTCAACCAT |
| Pink1-E4-1 | TATTAGTGAGTTTATAAATGTTTAAGGGGT | ACTAAATAAAAATACATAACTCTATTCTTCT |
| Pink1-E4-2 | AGAAGAATAGAGTTATGTATTTTTATTTAGT | AAAAAACTTCAACTCTTCTAAAACCAAAATTAC |
| Pink1-E5-1 | TGAGATAAAGTTTTATGTAGTTTAGGTTGGT | ACACACTACATAAAATAAAATAATATACAATAAAA |
| Pink1-E5-2 | TTTTATTGTATATTATTTTATTTTATGTAGTGTGT | TAACCTCTCTATCCCATCATTAACTACCTCCAC |

#### Table S5. Primer sequences for ChIP-qPCR.

| Name | Forward (5'-3') | Reverse (5'-3') |
|---|---|---|
| Pink1-E1/E1_a | GCTGGCTGCTACCACTCCTG | TCTGTCCCTCTTGGCCCAGG |
| Pink1-E1_b | GTGGAGACATTGGGTCAAGC | ACCACTAAACTGGGACCAAG |
| Pink1-E2 | CCAAGTAGCTGAGATTCCAG | CTCTGACATTAGAGGTACAC |
| Pink1-E3 | AACCTCAGCAGAGTGCACAG | TGTGAGTTTGAGGCCAGCTG |
| Pink1-E4 | GCCACAGGAGTGAGACAATG | CTATCCTGATTCCCTGCTTG |
| Pink1-E5/E5_a | GCTTTATGCAGGCTAGGTAG | GAAGAGGCCACACATGGAAG |
| Pink1-E5_b | TCATGCAGTGTGCTCTATGC | AGTCATCTGAAATGAGCCAG |
| Background | AGTCAAGAACTGACTTAAGG | AGCCACAGAGAAGAGCCAAG |
| Gene desert | TCCTCCCCATCTGTGTCATC | GGATCCATCACCATCAATAACC |

**Tables S1-S2**

Table S1. Mass spectrometry results of WT MuSCs over-expressing either empty vector or *Pink1*.

Table S2. Mass spectrometry results of WT MuSCs over-expressing *Pink1* treated with or without BTZ.
